# Translation of numerous upstream open reading frames rather than their products is essential for proliferation of human cells of distinct origin

**DOI:** 10.1101/2025.07.07.663426

**Authors:** Nikita M Shepelev, Elizaveta A Razumova, Alexandr I Lavrov, Stephen J Kiniry, Alexandr M Makaryuk, Renata D Bibisheva, Olga A Dontsova, Pavel V Baranov, Maria P Rubtsova

## Abstract

Upstream open reading frames (uORFs) are a widespread class of translated regions (translons) occurring in 5′ leaders of mRNAs and serving critical roles in post-transcriptional regulation. However, their specific biological activities in human cells remains to be fully elucidated. Here, we conducted a genome-wide CRISPR-Cas9 loss-of-function screen of 978 uORFs identified with ribosome profiling, across human cell lines of distinct origin (HAP1, A549 and HEK293T). A total of 155 uORFs were identified as being essential for cell proliferation. These uORFs showed a high cell-type specificity, with only a few being universally essential. Subsequent analysis has revealed that the primary reason underlying the uORF essentiality is not encoded micropeptides, but rather *cis*-regulatory mechanisms. Moreover, uORFs located within short 5′ UTRs were disproportionately sensitive to frameshift-inducing indels, which frequently lead to the uORF extension and overlap with the coding region (CDS), resulting in translational repression. Finally, by intersecting regions of essential uORF with ClinVar and dbSNP datasets, we identified naturally occurring variants with the potential to disrupt their function and contribute to disease phenotypes. These findings highlight a pervasive and underappreciated layer of translational control in human cells and establish uORFs as critical *cis*-regulatory elements with potential relevance to human health.

## Introduction

Any nucleotide sequence contains a large amount of short open reading frames (short ORFs) due to the random occurrence of start and stop codons in nucleotide sequences. Long ORFs are rare in random nucleotide sequences, yet they are typical in the sequences of RNA molecules coding for proteins due to a negative evolutionary pressure acting on stop codons within protein coding sequences (CDS). Therefore, the lack of stop codons within a single reading frame provides the evolutionary evidence for the existence of protein coding information (1). Hence, for a while, short ORFs have not been considered as potentially protein coding with a length of 100 codons being used as the lower CDS limit (2–4).

Although the translation of short ORFs occurring upstream of CDS (uORFs) have been known to play critical roles in the regulation of CDS translation in eukaryotes since the late 1980s – early 1990s (5–7), they have not been a subject of genome annotation, which has focused on the identification of encoded proteins rather than on the annotation of all translated regions (translons) (8). The advent of ribosome profiling revolutionized our understanding of genome translation by providing a technology for experimental identification of all translons independent of their length. In addition to CDS, numerous additional translons have been identified. Ribosome footprints have been found to map to long non-coding RNAs (lncRNAs) as well as to 5’ leaders and 3’ trailers of mRNAs that have been traditionally considered as untranslated regions (UTRs) (9–13). Furthermore, ribosome profiling data exhibit a distinct three-nucleotide periodicity, reflecting the stepwise movement of the 80S ribosome during translation (14) and showing a framing bias towards the translated reading frame. Utilization of this bias led to identification of translons overlapping or nested within annotated CDS (15).

The translation products of numerous translons, which were detected via ribosome profiling, were observed through mass spectrometry analysis of peptides presented by the major histocompatibility complex I (MHC I) (16–20). Recent studies suggest that many stable polypeptides, which associate with protein complexes, originate from short translons mapping to mRNAs leaders and non-coding RNAs (21).

Despite the widespread presence of translons lacking annotation as CDS (enigmatic translons), only a small fraction has been characterized in detail to elucidate their biological activity in cellular processes. Several studies have addressed the problem using CRISPR-Cas9 screening (22–25) and mass spectrometry (16, 18, 20, 26, 27). They have validated several examples of biologically relevant enigmatic translons. To help researchers with the task of understanding the biological significance of enigmatic translons, GENCODE has started to catalogue these elements identified with ribosome profiling (28).

Determining biological roles of enigmatic translons by sequence comparison remains challenging due to their short length and the evolutionary novelty of many examples (29). This complexity is further exacerbated by the frequent use of non-AUG start codons for translation initiation (30). Several large-scale studies have identified non-AUG initiated proteoforms (31–34) many of which are conserved (34, 35), and few mammalian examples have been well-characterized long time ago (36–38).

A noteworthy example is the *EIF4G2* gene, which encodes the translation initiation factor eIF4G2 (DAP5) without an AUG start codon (38). Additionally, CUG-initiated alternative proteoforms of cancer-related genes such as *MYC* and *FGF2* exhibit distinct functional properties (36, 37, 39, 40). Several non-AUG N-terminal extended variants of PTEN (31) play roles in regulating PI3K signaling, pre-rRNA synthesis, and cell proliferation (41–43). Moreover, ribosome profiling and proteomic studies suggest that the majority of uORFs are translated from non-AUG codons (44, 45). Some non-AUG starts are very efficient, reaching up to 70% of that of AUG in the optimal Kozak context and under stimulation of an RNA stem loop, as observed at the CUG start of the POLGARF protein encoded in the *POLG* mRNA (46).

uORFs represent a distinct class of regulatory translons located within the 5’ leader sequences of most mRNAs, since a half of all mRNAs contains at least one AUG upstream of CDS (47, 48). The exact biological roles of uORFs are difficult to characterize because one need to distinct between activity of their products and their effect on the translation of the downstream CDS (49, 50), since any change in the uORF sequence could potentially affect both facets of this class of translons. While in general, uORFs mildly suppress the translation of CDS, there are many examples of uORFs that play a dynamic regulatory role, allowing cells to modulate translation in a response to changing conditions (51–56). Moreover, non-AUG initiated uORFs introduce another layer of the regulation due to their variable and context-dependent translation efficiency (57, 58).

The most extensively studied mammalian regulatory uORFs are found in the *ATF4* gene, which encodes a master transcription factor of the integrated stress response (59). It was initially proposed that under stress conditions, the *ATF4* translation is induced through a ribosome re-initiation mechanism involving two AUG uORFs in its 5’ leader sequence (60). However, subsequent studies have revealed that this regulatory mechanism is considerably more sophisticated, involving a conserved stem-loop, a near-cognate CUG, and the process of nonsense-mediated decay, thereby highlighting the complex nature of the *ATF4*’s translational control (61, 62).

Another emerging area of interest is uORF-coded microproteins. For example, the ASDURF microprotein encoded by the *ASNSD1* mRNA uORF, is involved in a prefoldin-like chaperone complex that promotes medulloblastoma cell survival (24, 63). The *MIEF1* microprotein, L0R8F8 (a.k.a. AltMIEF1, MIURF, MIEF1-MP) regulates the assembly of mitoribosome by preventing formation of the full ribosome complex prior to the proper assembly of the large subunit (64, 65). In addition, an uORF in the *Protein Kinase C-eta* (*PKC-η*) mRNA encodes a peptide (uPEP2) containing a PKC pseudosubstrate motif, which inhibits the kinase activity of PKC family members (66).

In this study, we aimed to conduct a comprehensive analysis of human uORFs and to identify those that are essential for cell proliferation. We integrated ribosome profiling data with CRISPR-Cas9 screenings to systematically select essential uORFs across three distinct human cell lines. Our findings suggest that uORFs usually exert their function not by encoding microproteins, but by regulating the translation of the downstream CDS. Moreover, obtained results indicate that genes containing uORFs in short 5’ UTRs are more vulnerable to the translational repression due to frameshift generating indels within the uORF. These findings enhance our understanding of the uORF-mediated gene regulation and highlight potential candidates for further functional studies.

## Materials and methods

### Analysis and visualization of ribosome profiling data

uORFs were selected using Trips-Viz translated ORF detection tool (https://trips.ucc.ie/homo_sapiens/Gencode_v25/orf_translation/) using the following studies: Ji et al. 2016 (BJ and MCF10A) (67), Gameiro et al. 2018 (MCF10A) (68), Xu et al. 2016 (fibroblasts) (69), Zhang et al. 2017 (HEK293) (70), Calviello et al. 2016 (HEK293) (71), Iwasaki et al. 2016 (HEK293) (72), Park et al. 2016 (HeLa) (73), Crappe et al. 2015 (HCT116) (74), Fijalkowska et al. 2017 (HCT116) (75), Goodarzi et al. 2016 (MDA-MB-231) (76), Guo et al. 2014 (U2OS) (77), Wolfe et al. 2014 (KOPT-K1) (78), Werner et al. 2015 (hESC) (79).

Subcodon profiles of the highest-ranking results were then visually inspected and uORFs selected manually. Additional uORFs were taken from supplementary table 4 (Phase I Ribo-seq ORFs) from (28).

Aggregate ribosome profiling data were visualized in Trips-Viz browser selecting datasets used for uORF prediction. To visualize the data with the ribosome stalled at the initiation stage by harringtonine and lactidomycin, suitable data sets were selected from the following works: Ji16 (BJ), Zhang17 (HEK293), Fijalkowska17 (HCT116), SternGinossar12 (Human fibroblasts), BerkovichKinori16 (A549), Chen20 (iPSC, Cardiomyocytes) (PMIDS: 26900662, 29170441, 28541577, 23180859, 23180859, 32139545).

### uORF conservation analysis

The phyloP tracks were obtained from the UCSC Genome Browser the following genome alignments: a 30-way genome alignment (predominantly primates), a 241-way genome alignment (mammals), and a 100-way genome alignment (vertebrates). The PhyloCSF tracks downloaded from https://data.broadinstitute.org/compbio1/PhyloCSFtracks/ for 58-way genome alignment (mammals), and a 100-way genome alignment (vertebrates). The mean nucleotide (phyloP) or codon (PhyloCSF) scores of uORFs were calculated. Scores for start and stop codons were excluded from the analysis for PhyloCSF. Notably, in certain cases, some uORFs were not fully covered by scores due to the presence of gaps in the alignments and codons spitted between genomic segments of uORFs for PhyloCSF.

The *TRAM1* uORF consensus was obtained using the uORF4u tool (version 0.9.5) (80) under the following conditions: -an NP_055109.1 -bdb refseq_protein -bh 200 -bid 0.5 -mna 3 -ul 1000 -dl 300 -pc 0.3 -c eukaryotes. The consensus is based on 123 sequences.

### CRISPR/Cas9 library design and cloning

Guide RNAs (gRNAs) were selected with CHOPCHOP (81), uORFs required to have at least 2 valid gRNAs before being included. Next guide RNAs with the highest off-target activity were removed. The cut site distances of gRNAs to the nearest CDS or exon boundary were determined using GENCODE Release 25 basic annotation for transcripts with “basic” or “CCDS” tags.

The oligonucleotide pool was designed to have homologous ends matching those of the Esp3I-digested vector lentiGuide-Puro (Addgene #52963) and synthesized by Twist Bioscience. The oligonucleotide pool was amplified within the log-linear range of the reaction and cloned into lentiGuide-Puro, digested by Esp3I (Thermo Fisher), using NEBuilder HiFi DNA Assembly Master Mix (NEB). After ligation, the DNA was purified using isopropanol precipitation and electroporated into DH5α, resulting in an approximate 300x number of colonies per gRNA. The DH5α colonies were then scraped and the plasmid library was purified using EndoFree Plasmid Maxi Kit (Qiagen). To test library quality, it was amplified using Q5 polymerase and deep sequenced on Illumina HiSeq at 300x coverage (reads per gRNA).

### Cell culturing

The HAP1 cells were obtained from Horizon Discovery, the HEK293T and A549 cells were obtained from American Type Culture Collection (ATCC). The HAP1 cells and their derivative cell lines were maintained in IMDM (Servicebio) supplemented with 10% fetal bovine serum (FBS, HiMedia), 2 mM L-alanyl-L-glutamine (HiMedia), 100 units/ml penicillin, and 100 μg/ml streptomycin (HiMedia) in 5% CO_2_ incubator at 37°C. The HEK293T and A549 cells and their derivative cell lines were maintained in DMEM/F-12 (Servicebio) with the same supplementation. All cells were routinely tested for the absence of mycoplasma using MycoReport (Evrogen).

### gRNA library packaging and titer

The gRNA library was packaged into a lentiviral library using the transfection of the HEK293T cells in two T175 flasks with Lipofectamine 3000 (Invitrogen) according to manufacturer’s application note with a minor modification. In brief, 22.1 million HEK293T cells per T175 flask were incubated overnight in lentiviral packaging media (Opti-MEM I Reduced Serum Medium supplemented with GlutaMAX (Gibco), 5% FBS (HiMedia), and 1 mM Sodium Pyruvate (Gibco). Following day, the cells in two T175 flasks were transfected using 129 µl Lipofectamine 3000 reagent, 32 µg psPAX2 plasmid (Addgene #12260), 32 µg plasmid library, 6.3 µg pCMV-VSV-G plasmid (Addgene #8454), and 139 µl P3000. After 6 hours, the media was replaced with fresh lentiviral packaging media. Viral supernatant was collected 24- and 52-hours post-transfection and filtered through 0.45 µm PES filter. Optimal infection conditions were determined to achieve ∼30% of infected cells for each cell line. To do this, cells were infected with different volumes of viral supernatant in the complete media (see Cell culturing) at final concentration of 10 µg/ml polybrene (Sigma). 24 hours after infection, the cells were trypsinized and split into two wells of a 6-well plate in the complete media. One well out of two was supplemented with the appropriate concentration of puromycin: 0.5 µg/ml for HAP1 and HEK293T, 1 µg/ml for A549. Three days after selection, the cells were counted to calculate infection efficiency by comparing the number of cells with and without puromycin selection.

### CRISPR/Cas9 screening

Cas9-expressing cell lines were obtained by infecting primary cell lines with lentiviruses packaged with the lentiCas9-Blast plasmid (Addgene #52962) at MOI ≤ 0.1, and then selected using 5 µg/ml blasticidin S (Gibco). The resulting cell lines were infected with lentiviruses containing the gRNA library at a ratio of 20-40% infected cells and 1000x coverage (infected cells per gRNA) in quadruplicate. Following day, the cells were trypsinized and seeded in the complete growth media (see Cell culturing) supplemented with the appropriate puromycin concentration: 1 µg/ml for HAP1 and HEK293T, 1.5 µg/ml for A549. Two days later, the control population was collected, and the cells were subcultured: every two days for HAP1 and HEK293T and every three days for A549. The final cell populations were harvested eight days later (or nine days later for the A549 cells), following the collection of the control population. Genomic DNA was isolated using spin columns (Lumiprobe), and the gRNA-encoding regions were amplified by PCR within the log-linear range of the reaction using Q5 polymerase (New England Biolabs) and deep sequenced on Illumina HiSeq at 500–1000x coverage (reads per gRNA).

### Data analysis of CRISPR/Cas9 screening

Sequencing data were processed using MaGeCK (82) to calculate gRNA counts. Fold-change and p-values were calculated using MLE and paired RRA algorithms included in MaGeCK. The MaGeCK MLE results were processed by MAGeCKFlute (83) to analyze pathway enrichment using Kyoto Encyclopedia of Genes and Genomes (KEGG), Reactome, and Gene Ontology Biological Processes (GOBP) and compare to DepMap data (84).

### Cloning

The plasmid used to validate the screening results in a competition assay (LeGO-gG2) was previously made using LeGO-iG2 (Addgene #27341) and lentiGuide-Puro vector (Addgene #52963) (85). Individual guide RNA sequences were cloned to the gRNA scaffold by annealing and ligating the oligonucleotides (see Supplement Table S6) into Esp3I-digested vector.

The plasmid for the dual luciferase assay (pGL3-EFS-Rluc-v2), which contains firefly (Fluc) and *Renilla* (Rluc) luciferase under different promoters, was created from the pGL3-Rluc vector (a kind gift from Radik R. Shafikov, unpublished). The vector backbone was amplified using primers Fluc-v2-f and Fluc-v2-r (see Supplement Table S6), and the EF-1α core promoter was amplified from the lentiCas9-Blast plasmid (Addgene #52962) using primers EFS_fw and EFS_rev (see Supplement Table S6). These products were then assembled into the reporter vector using NEBuilder HiFi DNA Assembly Master Mix (NEB), introducing two Esp3I sites directly after the EF-1α core promoter and immediately before the Fluc ORF without a start codon.

The 5’ end of a transcript of interest, including the CDS codons, was amplified from cDNA obtained using the reverse transcriptase Magnus and oligo(dT)_15_ primer (Evrogen). The PCR-product and pGL3-EFS-Rluc-v2 were digested using Esp3I and ligated using T4 ligase (Evrogen). Frameshift mutants were obtained by overlap extension PCR from the wild-type sequences of the 5’ ends of transcripts (see Supplement Table S6).

### Transfection and virus production

HEK293T cells were transfected with plasmids in a 96-well format using the Lipofectamine 3000 Transfection Reagent kit (Invitrogen) according to the manufacturer’s protocol. The cells were harvested two days post-transfection for luciferase assays.

Lentiviral particles were produced using the Lipofectamine 3000 Transfection Reagent kit (Invitrogen) according to the manufacturer’s application note (see gRNA library packaging and titer), with reagents scaled down for a 6-well plate.

### Competition assay

Cells expressing Cas9 in the appropriate media (see Cell culturing) were transduced with viral particles in triplicate in a 12-well plate at a ratio of 40-60% infected cells. The percentage of EGFP-fluorescent cells was then assessed using flow cytometry on LongCyte (Challenbio) or SinoCyte (BioSino) over several days. The mean ratio of EGFP-positive cells to wild-type cells was then calculated for each well. The resulting values were then normalized by the ratio at day three following the transduction for each well. Thereafter, the means and standard deviations were calculated for each sample and respective timepoint.

### Luciferase assay

Luciferase activities were measured in accordance with the manufacturer’s protocol for TransDetect Double-Luciferase Reporter Assay Kit (TransGen Biotech), in a 96-well format, 2 days post-transfection. The signal was measured at regular intervals post-reagent addition using Victor X5 plate reader (Perkin Elmer).

### Statistical analysis

Statistical analysis was performed using GraphPad Prism 10.3.1 software (GraphPad Software). Statistical significance was determined using non-parametric tests, corrected for multiple comparisons if applicable.

## Results

### Identification on translated uORFs and analysis of their sequence conservation

A large amount of publicly available ribosome profiling datasets provides a valuable resource for identifying translated uORFs (86). To leverage these data, we analyzed aggregate ribosome profiling data and selected human uORFs using the Trips-Viz translated ORF detection tool (87) (Figure 1A). In order to maximize the number of potentially active uORFs, we included those with near-cognate start codons (differ from AUG by a single nucleotide). We considered an uORF to be a translon only if aligned ribosome footprints exhibited (i) strong framing bias matching that of the corresponding reading frame, (ii) an increase in read density at the start codon, and (iii) a decrease after the stop codon (Figure 1B). To expand this selection further, we also added translons from the “Phase I Ribo-seq ORFs” catalog (28), which includes only AUG uORFs (Figure 1A).

**Figure 1.**
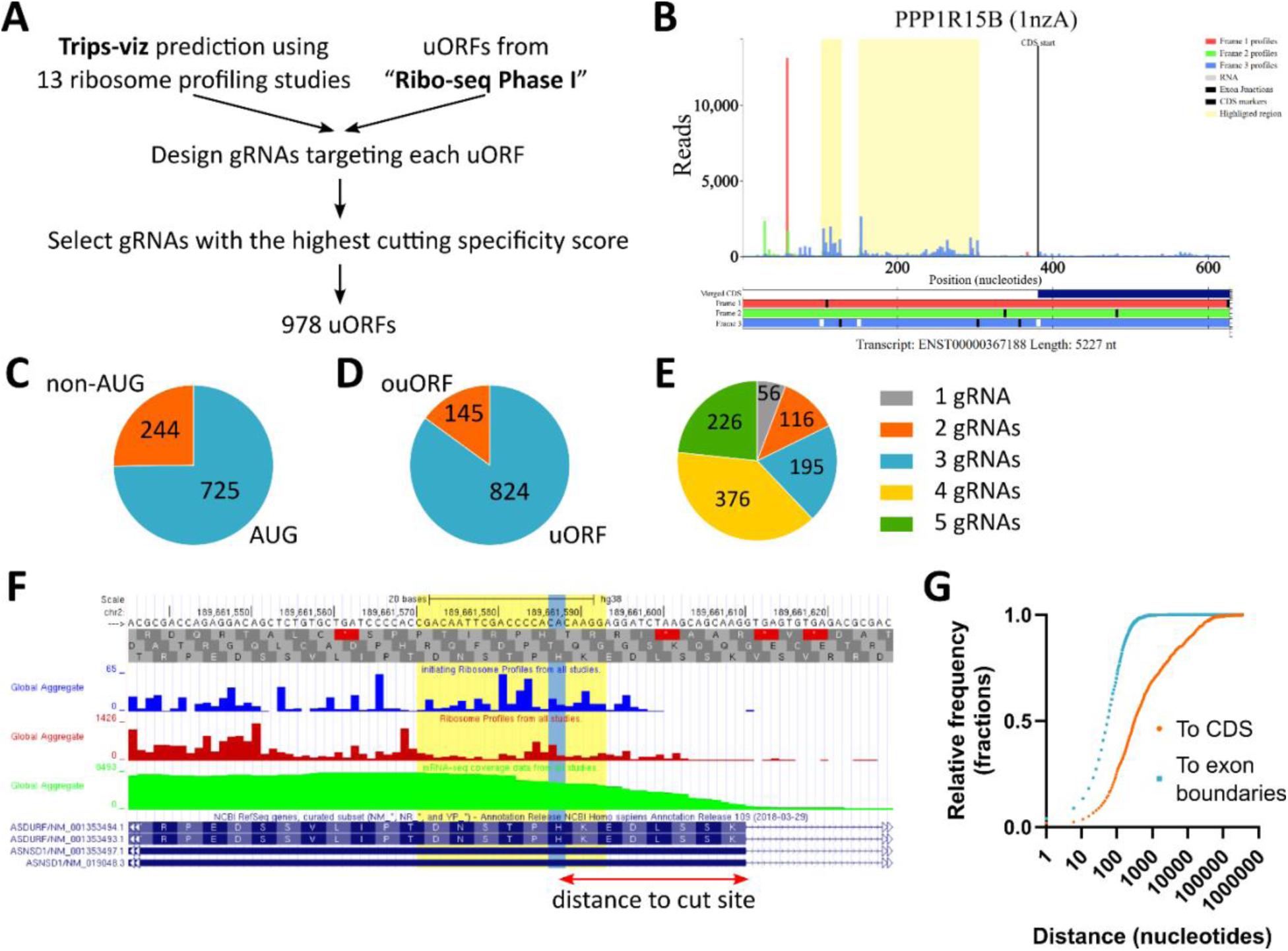
The selection of translated upstream open reading frames (uORFs) in human. (**A**) The uORF selection pipeline outlining the criteria used to pick uORFs. (**B**) An example of ribosome profiling data for the *PPP1R15B* mRNA, with selected uORFs highlighted in yellow. Translation hallmarks considered include: an increased ribosome read density at the start codon, the framing bias of three-nucleotide periodicity matching uORF reading frame, and a decrease in read density after the stop codon. (**C**) The distribution of start codons for selected uORFs, including AUG and near-cognate start codons. (**D**) Proportions of overlapping uORFs (ouORFs) and non-overlaping uORF (uORFs). (**E**) The distribution of gRNAs across uORFs, showing the number of gRNAs designed for each target. (**F**) The illustration of the gRNA selection in the GWIPS-viz browser (91). A targeted sequence and the PAM-motif are highlighted in yellow. The cut site is highlighted in blue. The distance to exon boundaries and coding regions (CDS) were considered. (**G**) The histogram showing distances of gRNA cut sites to the nearest exon boundary or CDS. The bin width is 5 nucleotides.

A widely employed functional genomics approach that identifies genetic elements influencing a specific phenotype is CRISPR screening. The simplest approach to identify functional translons is to assess cell proliferation after the translon disruption (assuming no activity unrelated to translation). Thus, we sought to systematically evaluate the uORF essentiality for cell proliferation. To this end, a guide RNA (gRNA) library for the CRISPR-Cas9 system was designed to induce indel mutations in the selected uORFs. Nevertheless, many uORFs were excluded due to technical constraints, such as the absence of a PAM-motif, the lack of specific and effective gRNAs, and the case of overlapping uORFs where all potential gRNAs result in a cut within the coding sequence (CDS). The selection of gRNAs for uORFs from Trips-Viz prediction and the “Phase I Ribo-seq ORFs” catalog resulted in a refined list of 969 targeted uORFs (Supplement Table S1), of which 725 (75%) are AUG uORFs (Figure 1C). The majority (824, 85%) of uORFs do not overlap with the CDS (Figure 1D). The length distribution of the selected uORFs exhibited a marked skewness towards shorter lengths, with a median length of 93 nucleotides (Supplement Figure S1A), consistent with previous reports (88, 89).

For the majority of uORFs (797, 82%), we succeeded in designing 3–5 gRNAs per uORF. However, for the remaining uORFs (172, 18%), we were only able to design 1–2 gRNAs due to the previously mentioned constraints (Figure 1E). In order to ensure both specificity and efficiency, selected gRNAs were optimized for the high on-target activity and minimal off-target effects. Furthermore, we refined the selection process by ensuring that gRNA cut sites were distant from exon boundaries and the CDS (Figure 1F). The majority of uORFs met these criteria except for rare cases of alternative isoforms (Figure 1G). The final gRNA library also included 102 non-targeting gRNAs (negative controls) and 56 gRNAs targeting 14 common essential genes involved in critical cellular processes (positive controls) (Supplement Table S2). In total, the library contained 3,665 gRNAs (Supplement Table S3).

We then evaluated the selected candidates by assessing their nucleotide conservation using uORF average phyloP scores from three comparative genomic alignments: a 30-way genome alignment (mostly primates), a 241-way genome alignment (mammals), and a 100-way genome alignment (vertebrates) (90) (Supplement Table S1). The rationale behind our selection of phyloP stems from the premise that functionally conserved uORFs may not necessarily require conserved peptides; instead, a nucleotide-level conservation may indicate their biological significance. Notably, 765 uORFs (79%) exhibited positive phyloP scores across all three alignments (Supplement Figure S1B). As anticipated, phyloP scores demonstrated significant correlations across the alignments (Supplement Figure S1С–E). Weak negative correlation was found between the uORF length and the average phyloP score, which was significant only for the 30- and 241-way alignments, indicating that nucleotide conservation of uORFs in our set is not specifically dependent on their length (Supplement Figure S1F–I). As expected, uORFs overlapping CDS (ouORFs) exhibited significantly higher levels of nucleotide conservation in all alignments due to their partial overlap with conserved CDS (Supplement Figure S1J). Additionally, phyloP scores for AUG- and non-AUG-initiated uORFs were comparable regardless of ORF type (uORF or ouORF) (Supplement Figure S1K–L).

### Many uORFs are essential for the proliferation of near haploid cells of leukemic origin

We selected near-haploid HAP1 cells (92) as the primary cell line in this study, as their minimal genomic complexity almost eliminate the impact of copy number variation on CRISPR/Cas9 screening results (93). To perform a CRISPR/Cas9 loss-of-function knockout screen, HAP1 cells were transduced overnight with lentiviruses containing the gRNA library at a multiplicity of infection (MOI) of 0.3 ensuring one gRNA in most cells. The cells were then selected with puromycin for 2 days, after which the control population was collected. The remaining cells were passaged for 8 days to collect the final population (Figure 2A). Quality control metrics confirmed high biological reproducibility and sample quality (Supplement Figures S2A–D). Strong depletion of positive control genes validated the robustness of the screening conditions. Additionally, strong enrichment of non-targeting gRNAs suggested that many uORF gRNAs were instead depleted. Thus, mutations in many uORFs inhibited cell proliferation (Figure 2B, Supplement Figure S2E).

**Figure 2.**
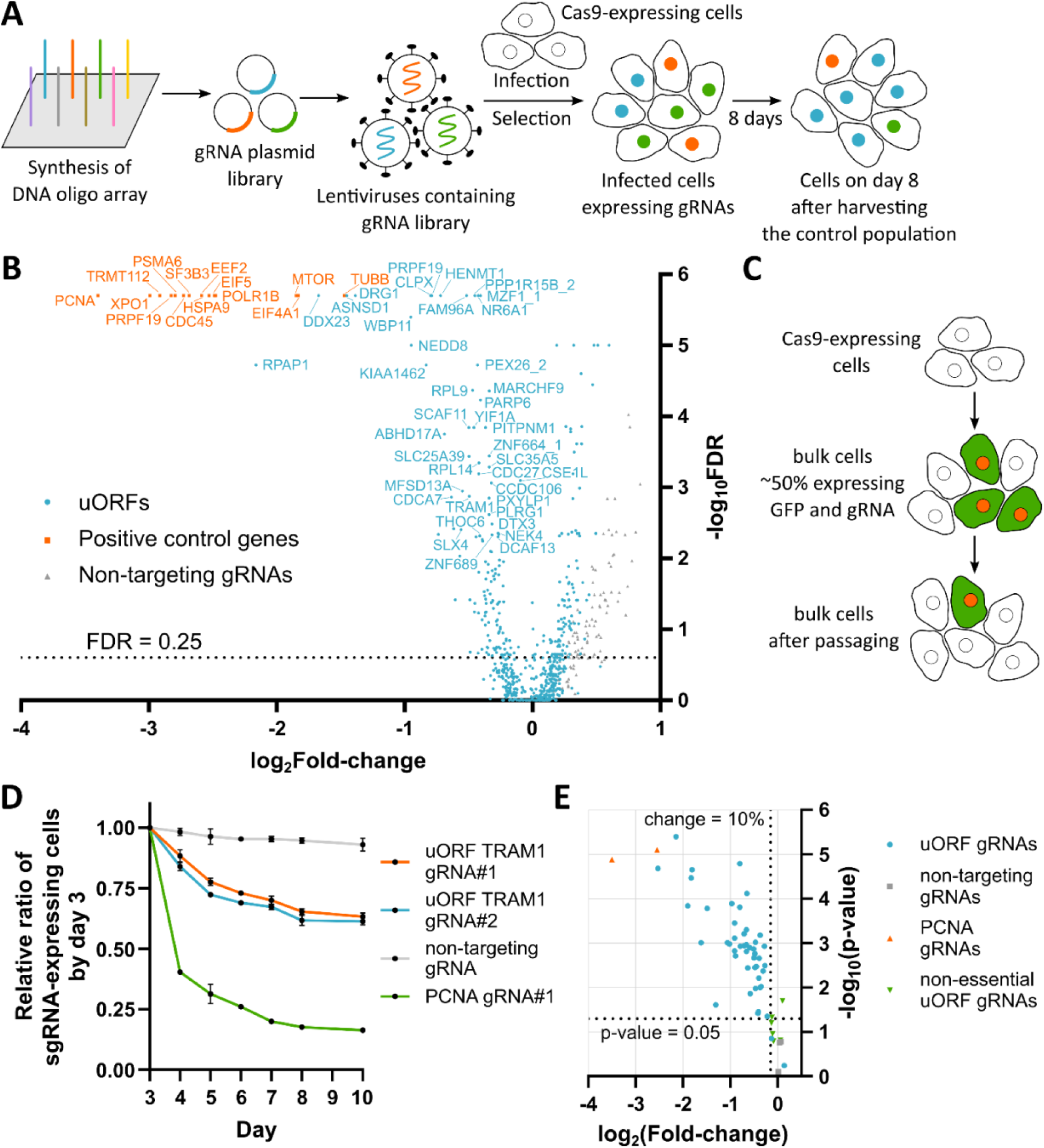
Numerous uORFs are essential for proliferation in HAP1 cells. (**A**) Schematic overview of the CRISPR-Cas9 loss-of-function screening workflow. (**B**) Volcano plot illustrating screening results in HAP1 cells. Each dot represents an uORF (blue), a positive control gene (orange), or a non-targeting guide RNA (grey). The top 40 uORFs with the strongest depletion are alongside the tested essential genes. (**C**) Schematic representation of the competition assay used for screening validation. HAP1 cells were transduced with lentiviruses expressing individual gRNAs and the green fluorescent protein (GFP), enabling the proportion of uORF-disrupted cells to be assessed. (**D**) Representative results of the competition assay. Lentiviral transduction with GFP and gRNA enables the monitoring of the proportion of uORF-disrupted cells over time. (**E**) Validation of selected uORFs using individual gRNAs in HAP1 cells. Fold changes in the fractions of GFP-positive cells were measured between day 3 and day 10 post-transduction, and p-values were calculated in order to assess the significance of the depletion.

In total, 135 uORF passed the significance threshold at the false discovery rate (FDR) of 0.25 (Figure 2B, Supplement Table S4). Among these, several previously characterized uORFs were found to be also essential for cell proliferation. For example, the uORF in the *ASNSD1* gene (now annotated as *ASDURF*) had a critical impact, comparable to the positive control genes essential for HAP1 survival (Figure 2B). Other significant hits included uORFs in *PPP1R15B* (94, 95), *TLNRD1* (96), *EIF2D* (97), *MIEF1* (65, 98). However, most of top hits had not been previously studied. Pathway enrichment analysis did not reveal significant enrichment for top hits, consistent with the idea that essential genes are usually involved in multiple pathways (99).

To further validate our findings, we tested the top uORFs using individual gRNAs in a competition assay (Figure 2C). HAP1 Cas9 cells were transduced with lentiviruses expressing both the gRNA and the green fluorescent protein (GFP), ensuring a roughly equal mixture of wild-type cells and GFP-positive uORF-disrupted cells. We then monitored the fraction of infected cells over several days post-transduction (Figure 2D). This validation confirmed the screening results for the majority of tested hits, while non-targeting control gRNAs had no effect on cell proliferation (Figure 2E). Additionally, several non-hit gRNAs also showed no effect, suggesting that the observed decrease in proliferation was not merely due to DNA damage. Moreover, the effects of uORF gRNAs do not correlate with the distance to the nearest CDS (Supplement Figure S2F) or exon boundary (Supplement Figure S2G), indicating no direct relationship between the gRNA effect and disrupted mRNA biosynthesis. The observed effects correlated well with the screening data, despite the substantial difference in experimental setups. This confirms the reliability of the screening results (Supplement Figure S2H).

An important question is what role of the identified targets is essential: regulatory or coding? Thus, to determine whether the observed effects were due to uORF disruption rather than subsequent CDS dysfunction, we compared our fold-change data with gene essentiality information from the DepMap database (84). For several genes, uORF knockout effects did not correspond to gene essentiality (Supplement Figure S2I), suggesting that mutations in these uORFs affect cell proliferation through their products rather than via a disruption of the CDS translation. Among them, *ASDURF* showed the strongest effect.

### uORFs essentiality is highly cell specific

Whole-genome CRISPR screenings across different cell lines have shown that gene essentiality can be either universal or highly cell-type specific (84). The same principle may apply to uORF function, given that uORFs typically serve regulatory roles, which may vary across different cellular contexts. HAP1 cells originate from chronic myeloid leukemia KBM7 cells and possess a unique genetic background (92). Therefore, we searched for essential uORFs in cell lines of different origins.

To this end, we performed CRISPR/Cas9 loss-of-function screening in both adenocarcinoma (A549) cells and non-cancerous human embryonic kidney (HEK293T) cells, following the same protocol as for HAP1 cells (Figure 2A). Quality control metrics confirmed high biological reproducibility and quality for both A549 and HEK293T samples (Supplement Figures S3A–D and S4A–D). While positive control genes demonstrated strong depletion in all cell lines, the effects were less pronounced in HEK293T cells (Figure 3A, Supplement Table S4) and A549 cells (Figure 3B, Supplement Table S4) compared to HAP1. As was previously observed, we noted a strong enrichment of non-targeting gRNAs, indicating that the knockout of numerous uORFs inhibited cell proliferation (Supplement Figures S3E and S4E). In accordance with the HAP1 screen, gene enrichment analysis did not reveal significant pathway enrichment, consistent with the assumption that essential genes act in multiple distinct pathways (99). In a similar manner, the effects of uORF gRNAs do not correlate with the distance to the nearest CDS (Supplement Figure S3F and S4F) or exon boundary (Supplement Figure S3G and S4G), indicating that there is no direct relationship between gRNA effect and disrupted mRNA biosynthesis.

**Figure 3.**
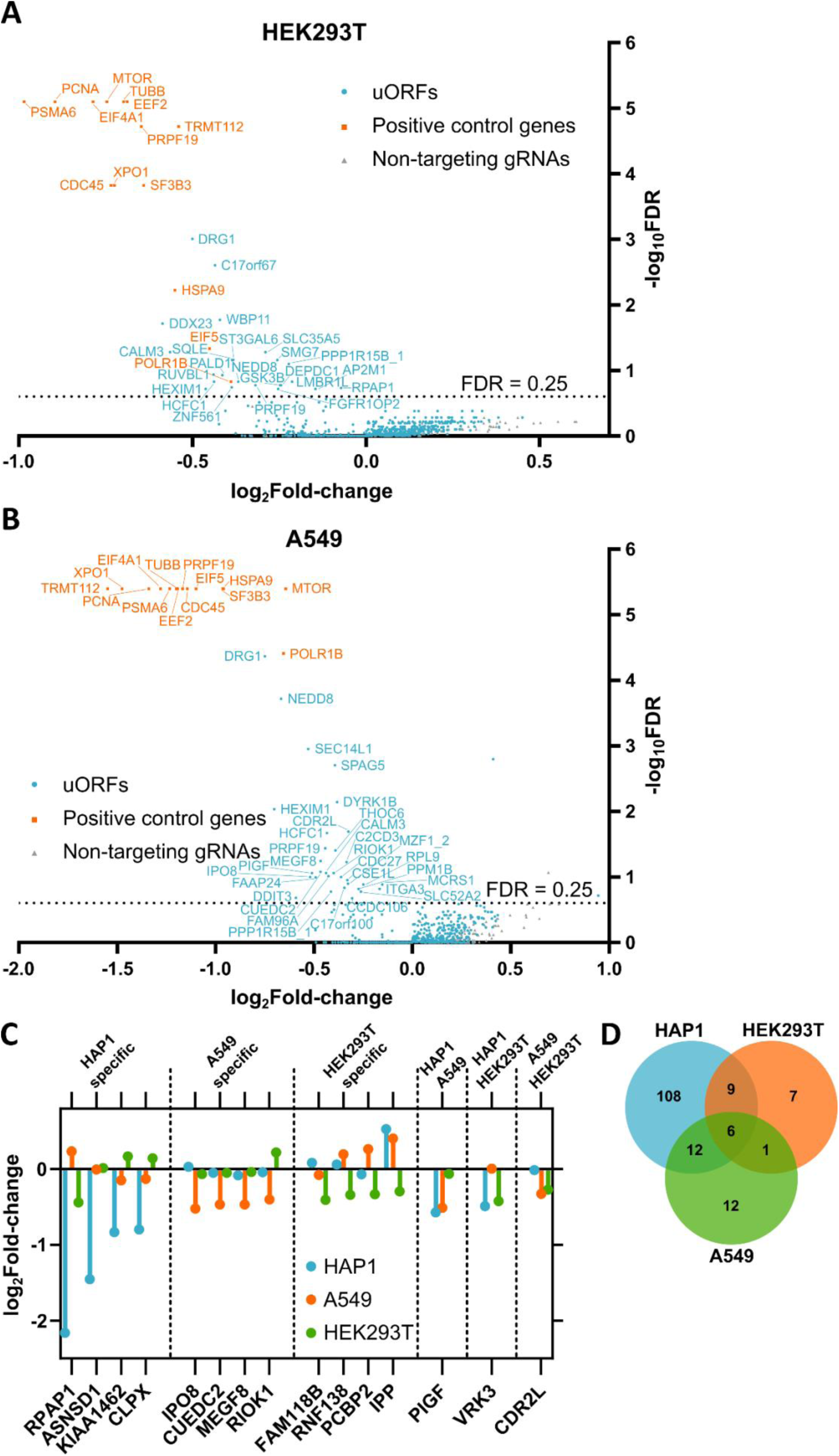
The majority of uORFs essentiality are cell-specific. (**A**) Volcano plot illustrating the screening results for HEK293T cells. Each dot represents an uORF (blue), a positive control gene (orange), or a non-targeting guide RNA (grey). The top 23 uORFs which passed the FDR threshold and exhibited the strongest depletion are labeled alongside the tested essential genes. (**B**) Volcano plot demonstrating screening results for A549 cells. Each dot represents an uORF (blue), a positive control gene (orange), or a non-targeting guide RNA (grey). The top 31 uORFs which passed the FDR threshold and exhibited the strongest depletion are labeled alongside the tested essential genes. (**C**) Log_2_Fold-changes of uORFs showing universal and specific essential uORFs. Some uORFs were depleted across two or three three cell lines, while others exhibited cell-type-specific effects. (**F**) Venn diagram illustrating the relationship between the results of three CRISPR screens for uORFs that passed the threshold (FDR < 0.25).

In total, 23 and 31 uORF passed the significance threshold (FDR < 0.25) in HEK293T and A549, respectively. Among these, several previously characterized uORFs were found to be essential for cell proliferation. In HEK293T cells, these included uORFs in *SPAG5* (100), *PRPF19* (85), *MZF1* (101), *PPP1R15B* (94, 95), and *DDIT3* (102). In A549 cells, the essential uORFs included those in *PRPF19* (85), *PPP1R15B* (94, 95), *SLC35A4* (95, 103). However, as in the HAP1 screening, most of the top hits had not been previously explored.

In order to validate our experimental conditions, we performed competition assays in A549 and HEK293T cells. Cas9-expressing A549 or HEK293T cells were transduced with lentiviruses expressing both gRNA and the GFP protein, ensuring a roughly equal mix of wild-type cells and GFP-positive uORF-disrupted cells. The fraction of infected cells was monitored over the period of several days post-transduction. A gRNA targeting the *PCNA* gene confirmed the robustness of screening conditions, while a non-targeting control gRNA had no effect on cell proliferation (Supplement Figures S3H and S4H). Interestingly, the observed effect of the *PCNA* gRNA in HEK293T cells was less pronounced compared to A549 cells, aligning with the weaker screening effects in HEK293T compared to HAP1 and A549 cells (Figure 3A). Nevertheless, we confirmed the gRNA effects for *CSK3B* and *SLC35A5* uORFs in HEK293T cells (Supplement Figure S3H). In A549 cells, validation by the competition assay was complicated by low resolution between GFP-expressing and wild-type cells.

Subsequently, we compared the fold-change values from our screening data to the gene essentialities from the DepMap database. In both HEK293T cells (Supplement Figures S3I) and A549 (Supplement Figures S4I), many uORF knockout effects did not correlate with gene essentiality, suggesting that these uORFs may have independent functions, rather than regulating CDS translation.

To investigate cell-type specificity, we compared uORF screening results across HAP1, A549, and HEK293T. The overall uORF essentiality profiles exhibited substantial variability between the cell lines, demonstrating low correlation with each other (Supplement Figures S5A, S5C, S5E). The effects of gRNAs targeting uORFs exhibited slightly higher correlations between all cell lines (Supplement Figures S5B, S5D, S5F). Even for commonly essential genes, the observed effects correlated moderately, implying that gene dependencies vary considerably depending on cellular origin. As anticipated, non-targeting control gRNAs exhibited no correlation between cell lines (Supplement Figures S5B, S5D, S5F). However, while most uORFs displayed high cell-specific essentiality, some uORFs were depleted in a similar manner across all three cell lines, whereas others exhibited completely different effects depending on the cell type (Figure 3E). In total, 155 essential uORFs were identified across all three cell lines; of these, only 6 were shared among them (*HCFC1*, *HEXIM1*, *CALM3*, *NEDD8*, *PRPF19*, *DRG1*) (Figure 3F).

We then compared our results with those of previous CRISPR-Cas9 screenings of newly identified translons, commonly termed “non-canonical ORFs” (22, 23, 25). Even a direct comparison of targeted uORFs is challenging given that the studies use different transcriptome annotations and approaches for translons selection. Furthermore, a comparison of the effects across studies was challenging due to differences in statistical methodologies. Therefore, firstly we simply compared the genes that encode translated short ORFs. The gene set in the present study shares approximately a third of its genes with Chen et al. (22), approximately a half with Schlesinger et al. (25), and strongly deviates from Prensner et al. (23) (Supplement Figure S6A). It is noteworthy that approximately one-third of the genes were examined for the first time in this study. In order to circumvent limitations imposed by statistical methodologies, we analyzed shared gRNAs used in other studies. Specifically, 165 identical gRNAs were identified from Chen et al., 14 from Prensner et al., and 102 from Schlesinger et al. However, a small number of identical gRNAs in Prensner et al. cannot provide reliable comparison. Therefore, we proceeded to examine the other studies. No correlation was observed between the gRNA effects in iPSC cells from Chen et al. and cells in the present study. We found a weak correlation between K562 from Chen et al. and cells in the present study, with the correlation being slightly higher for the K562 and HAP1 pair likely due to their shared leukemic origin (Supplement Figure S6B–G). Correlation coefficients were on average higher for 9 pairs produced by comparing three cell lines in the present study and cells from Schlesinger et al (Supplement Figure S7). Notably, these correlation levels were comparable to those between HAP1, A549, and HEK293T in the present study, suggesting a high dependency of uORF essentiality on cell context.

### Regulatory activity of most uORFs, rather than peptide-coding capacity, is crucial for cell proliferation

A total of 155 essential uORFs were identified, out of 969 tested (16%), across three cell lines. The distinguishing features of these uORFs were then analyzed in comparison to the rest. The selected uORFs did not significantly differ from non-hit uORFs in terms of start codon usage, type, or source (Supplement Figure S8A–C). Subsequently, we calculated start codon context scores using available data (104, 105) and observed no significant difference between the two groups (Supplement Figure S8D). Nevertheless, non-AUG context scores are less reliable by their nature, therefore we compared the scores for AUG uORF only. Screening hits exhibited significantly higher context scores compared to non-hits (Supplement Figure S8E). Indeed, the disruption of more active translated uORFs should result in more pronounced effects on cell proliferation regardless of their function. In addition, we unexpectedly discovered that essential uORFs tended to be shorter (Supplement Figure S8F) and be located in shorter 5’ UTRs (Supplement Figure S8G), while unimportant parameters such as the uORF GC content (Supplement Figure S8H) and the CDS length (Supplement Figure S8I) were not different between the two groups. In order to achieve a more precise understanding of the underlying reasons, we conducted a subsequent analysis of the uORF position within transcripts. The essential uORFs have a start codon located significantly closer to the 5’ end (Supplement Figure S8J). This tendency persisted for relative start codon positions within 5’ UTR (Supplement Figure S8K). The positions of uORF stop codons followed the same principle for exact numerical values (Supplement Figure S8L) as well as for normalized values relative to the 5’ UTR length (Supplement Figure S8M). It is noteworthy that the distances between CDS and uORF were slightly shorter for essential uORF (Supplement Figure S8N). In conclusion, the most important parameters of uORFs that demonstrated a negative association with cell proliferation in the screening were the start codon in a better context, shorter length and location closer to the 5’ end.

Next, we examined the evolutionary conservation of these essential uORFs at the nucleotide level. Contrary to expectations, PhyloP score analysis revealed no significant difference between the median scores of non-essential uORFs and essential uORFs across 30-, 100-, and 241-way genome alignments (Supplement Figure S8O). Given the lack of enrichment for conserved uORFs as conserved nucleotide sequences, we investigated whether the essential uORFs are enriched with microprotein-coding uORFs, given that our dataset includes examples of known microproteins.

Among these, ASDURF stands out as a well-characterized microprotein with a known function. The validation of the screening results in HAP1 cells using individual gRNAs in a competition assay demonstrated that indel mutations in the *ASNSD1* uORF significantly reduced cell proliferation, whereas mutations within the CDS had a much weaker effect (Supplement Figure S9A). These findings align with a recent report highlighting the essential role of ASDURF in the medulloblastoma context (24). Another microprotein of known function, L0R8F8 (a.k.a. AltMIEF1, MIURF, MIEF1-MP) was also identified in our dataset.

In order to identify additional potential microproteins, we analyzed protein coding potential of uORFs using PhyloCSF data for sequence alignments of 58 mammals or 100 vertebrates. The PhyloCSF scores for all uORF codons except for start and stop codons were then averaged. The results predicted that only 49 uORFs (5.0%) and 22 uORFs (2.2%) are under considerable purifying selection (above zero scores) in 58 mammals and 100 vertebrates, respectively, a proportion consistent with previous reports (106). As anticipated, the PhyloCSF scores were significantly lower for the vertebrate alignment (Supplement Figure S9B). Notably, we observed no bias in protein coding potential for ouORFs in contrast to nucleotide conservation (Supplement Figure S9C). Non-AUG uORFs have slightly elevated PhyloCSF scores, suggesting the conservation of certain non-AUG uORFs (Supplement Figure S9D). It is important to note that some uORFs can employ specific peptides for their regulatory activity, thus such cases may also display elevated “protein coding potential” (51, 55). In the list of essential uORFs, solely 8 and 4 uORFs exhibited PhyloCSF scores above zero for sequence alignments of 58 mammals and 100 vertebrates, respectively. Furthermore, a comparison of essential and non-essential uORFs revealed no significant difference in their average PhyloCSF scores, indicating that protein coding potential is not the primary determinant of uORF essentiality (Supplement Figure S9E). However, it is possible that our list contains some evolutionarily young microproteins, which have strongly negative PhyloCSF scores as a result of biases introduced by large sequence datasets, but nevertheless possess functional importance (107). Additionally, if a translon is highly conserved the PhyloCSF analysis will result in near zero score. These cases can be distinguished by very high phyloP scores.

In order to identify additional functional microprotein-encoding uORFs, we adopted an alternative approach. This involved searching for essential uORFs within genes with a low number of dependent cell lines in the DepMap database. The rationale behind this strategy is that, if the CDS of a gene is non-essential but its uORF is essential, this suggests that the uORF encodes a functional peptide. For instance, the *ASNSD1* uORF (ASDURF) exhibits a clear compliance with this principle (Supplement Figure S9A). One such candidate was *TRAM1*, a gene for which no significant dependency in any cell line, including HAP1 (with near zero effects) (Supplement Figure S2I), according to the DepMap data. Ribosome profiling suggests active translation of the *TRAM1* uORF within the first exon, and profiling of ribosomes arrested at the initiation stage also supports translation initiation at the uORF start codon (Supplement Figure S10A). Notably, this uORF exhibits significant conservation across mammalian species (Supplement Figure S10B). Moreover, HLA peptidomics has identified a single unique peptide, thereby suggesting the presence of the uORF-encoded peptide within human cells (Supplement Figure S10B).

In order to validate this finding, we designed a series of gRNAs targeting its promoter, uORF, 5’ UTRs upstream and downstream of uORF, and CDS in the first exon and downstream (Supplement Figure S10C). Surprisingly, competition assays showed that mutations within the *TRAM1* CDS affected cell proliferation of HAP1 cells, contradicting DepMap data (Supplement Figures S10C–D). Furthermore, the 5’ UTR of *TRAM1*, including uORF, was also found to be crucial for cell proliferation, thereby indicating the significant role of the entire 5’ UTR in the *TRAM1* expression (Supplement Figures S10C–D). These results suggest that *TRAM1* plays essential roles in some cellular contexts, for example HAP1.

Taken together, these findings indicate that, overall, the essentiality of most identified uORFs may be driven by their regulatory activity rather than their peptide-coding capacity. Despite this, we cannot exclude the potential protein-coding capacity of some selected uORFs (Supplement Figure S2I, S3I, and S4I). In order to explore regulatory activity of uORFs, we examined the effects of frameshift mutations in several essential uORFs using a luciferase reporter. We cloned the 5’ end of transcript of interest, including the CDS codons, upstream of the firefly luciferase (Fluc) ORF, which lacks a start codon, acts as the CDS, and is regulated by the EF1α promoter. *Renilla* luciferase (Rluc) is expressed from the SV40 promoter and serves as an internal control (Figure 4A). Manual inspection of 5’ UTR sequences indicated that indels frequently extend the uORF, thereby creating an overlap with the CDS. Another, albeit less prevalent, possibility is the occurrence of an uORF-CDS fusion. Indeed, a point insertion or deletion in the *SLC25A39* uORF resulted in an overlap between CDS and uORF in both cases, leading to significant repression of the CDS translation due to the impossibility of reinitiation (Figure 4B). Point indels in *YIF1A* result in divergent outcomes. The deletion of a single nucleotide leads to the long overlap between the CDS and uORF (Figure 4C), and a point insertion results in an uORF-CDS fusion. The overlap in *YIF1A* also ultimately result in downregulation of the CDS (Figure 4C). Nevertheless, the fusion creates the N-terminal protein extension that may increase the CDS translation in some cases (97), but which may be toxic to the cell. Such fusion may compromise the stability or activity of the fusion protein despite the absence of translational repression.

**Figure 4.**
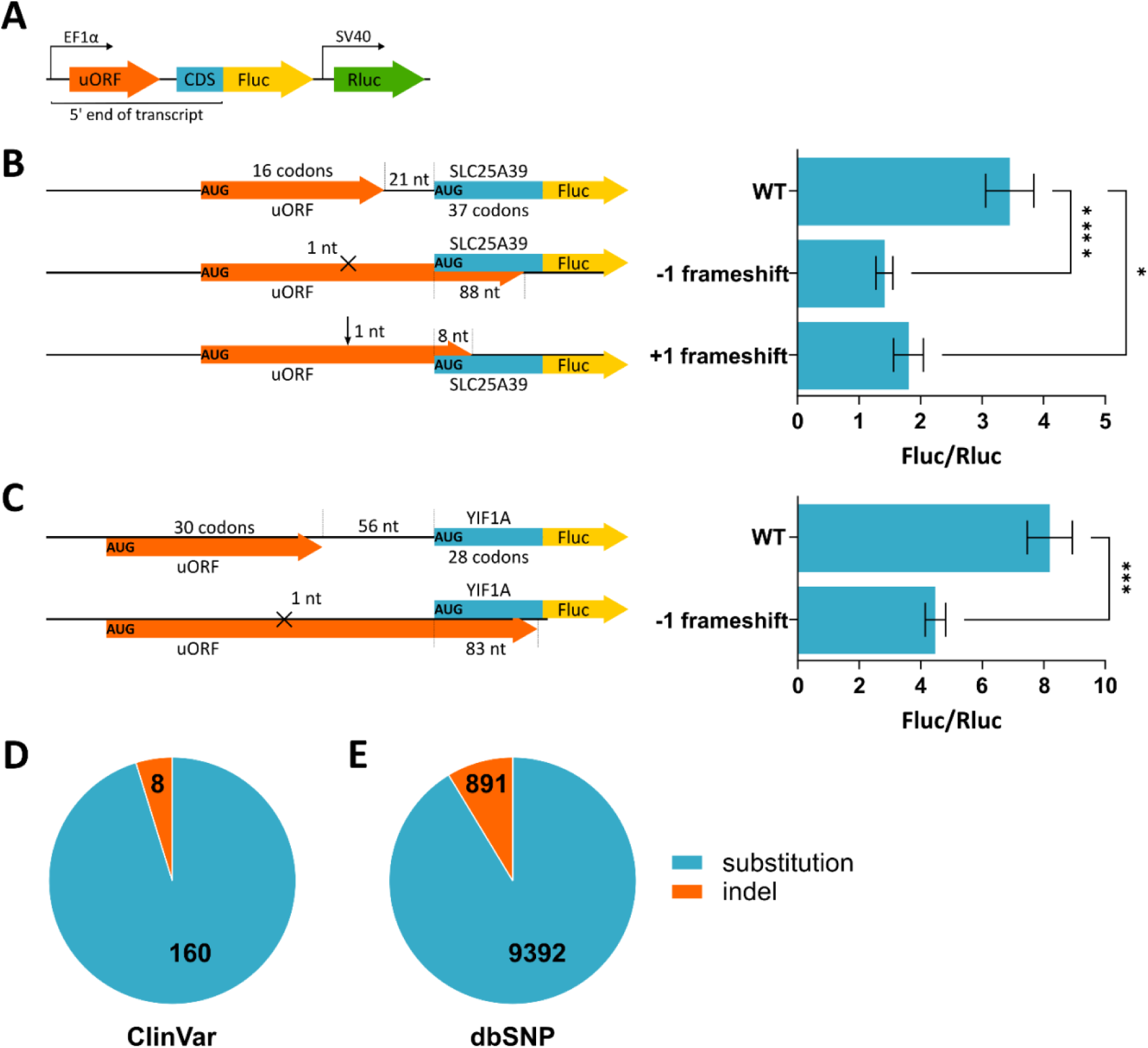
Indels in uORFs often lead to the repression of CDS translation. (**A**) Schematic representation of the plasmid reporter used to evaluate the impact of uORFs on the CDS translation. (**B**) Schemes of *SLC25A39* reporter constructs showing the outcome point mutations. Mean relative luciferase activities ± SD after transfection with the indicated constructs are shown (right). The Kruskal–Wallis test was conducted, followed by the Dunn’s multiple comparisons test. * – adjusted p-value < 0.05, **** – adjusted p-value < 0.0001. (**C**) Schemes of *YIF1A* reporter constructs showing the outcome point mutations. Mean relative luciferase activities ± SD after transfection with the indicated constructs are shown (right). *** – p-value < 0.001, determined by the Mann-Whitney U test. (**D**) A number of known mutations in essential uORFs from ClinVar (**E**) A number of known mutations in essential uORFs from dbSNP.

Given that mutations in essential uORFs can impact the CDS translation, naturally occurring nucleotide variations within these uORFs could contribute to human diseases (108, 109). In order to investigate this possibility, we analyzed single-gene variants in the ClinVar and dbSNP databases. ClinVar data for 5’ UTR regions was limited, presumably due to the conventional focus on CDS mutations. In total, 168 variants were identified in 30 uORFs, with only 8 indels present, all of which affect the CDS also (Figure 4D). Among the 53 variants that did not overlap with the CDS and should affect only the uORF, 28 were reported as benign or likely benign, while 25 have uncertain significance or conflicting classifications of pathogenicity. Nevertheless, we identified insertions, deletions, and single-nucleotide variants that could disrupt uORF regulatory functions, particularly those leading to frameshifting mutations (Supplement Table S5). The dbSNP database contained a greater number of variants, resulting in uORF frameshifts or codon substitutions, which potentially affect uORF-mediated regulation (Figure 4, Supplement Table S5). Specifically, 10,283 variants were identified across all 155 selected uORFs, with 891 indels present (Figure 4E). Among these, 813 (in 141 uORFs) did not affect the CDS. Additionally, 9,232 substitution variants (across all 155 uORFs) did not overlap with the CDS, and thus should only solely impact uORF function.

## Discussion

Upstream open reading frames (uORFs) were discovered long before the sequencing of the human genome (110–112). However, their widespread presence in eukaryotic genes was only established with the advent of high-throughput techniques such as ribosome profiling (also known as ribosome footprinting) (9–13, 95). Recent advances in ribosome profiling and mass spectrometry have revealed the pervasive translation of not only uORFs but also many other transcriptome regions previously classified as “non-coding” (18–21). Despite their widespread distribution, the functional significance of uORFs remains largely underestimated.

Currently, most high-throughput and targeted studies focus on identifying novel translons, including uORFs, that encode microproteins. However, compared to the growing number of studies on microproteins, examples of uORF-encoded microproteins remain the exception rather than the rule. The majority of known microproteins coded by newly identified translons is associated with long non-coding RNAs (lncRNAs), even though novel translons identified with ribosome profiling data occur in similar numbers in both lncRNAs and mRNAs (predominantly in 5′ leaders) (28).

Among the well-characterized microprotein-coding uORFs are ASDURF (63), L0R8F8 (a.k.a. AltMIEF1, MIURF, MIEF1-MP) (95, 64, 65, 98), SLC35A4 (95, 103), uPEP2 (58). These microproteins have independent function or modulate the activity of the main CDS-encoded protein. However, most uORFs likely serve regulatory functions rather than encoding conserved peptides. Importantly, uORF often play critical regulatory roles that are not necessarily dependent on peptide sequence conservation. Instead, their positioning within the 5’ UTR and codon composition are key factors in their function.

In this study, we employed a CRISPR-Cas9 screening approach to identify human uORFs that are essential for cell proliferation. Instead of focusing on protein coding potential, we selected uORFs based on ribosome profiling data and filtered them by the availability of the most reliable gRNAs (Supplement Table S3). For the screening, we initially utilized near-haploid human HAP1 cells, the cell line that minimize the influence of copy number variation, a common confounding factor in many cancer cell lines.

To analyze the data, we applied a rigorous and unbiased approach using MaGeCK (82). While this conservative analysis may have underestimated hits with only one or two effective gRNAs, we observed that even single-gRNA hits could be successfully identified, demonstrating their potential relevance (Figure 2B, Supplement Table S4). However, such cases require further thorough validation.

Using HAP1 cells, we identified numerous undescribed essential uORFs, but these cells represent only a single cellular context. To address cell-type-specific differences and estimate a proportion of generally essential uORFs, we conducted screenings in two additional cell lines of distinct origins: adenocarcinomic A549 cells (Figure 3A, Supplement Table S4) and non-cancerous HEK293T cells (Figure 3B, Supplement Table S4). The number of significant uORF hits in these cell lines was lower than in HAP1 cells, likely due to the hypotriploidy as an influencing factor.

We then compared the effects of gRNAs across different cell lines and observed low correlation between all cell line pairs, with HEK293T results deviating the most (Supplement Figures S5B, S5D, S5F). Interestingly, even gRNAs targeting generally essential genes, which served as positive controls, showed only moderate correlation across cell lines, highlighting strong differences in their gene dependencies. As expected, non-targeting gRNAs exhibited no correlation between cell lines (Supplement Figures S5B, S5D, S5F). When averaging gRNA effects by uORFs, we observed lower correlation (Supplement Figures S5A, S5C, S5E), likely due to variations in gRNA efficiencies across cell lines. However, the overall correlation remained low, and only six uORFs were essential across all three cell lines. Thus, our CRISPR-Cas9 screening demonstrated that uORF essentiality is highly cell-type-specific, with only a handful of uORFs being universally essential for proliferation, including previously reported uORFs in *PPP1R15B* (94, 95) and *PRPF19* (85).

We then compared our findings with those of previous studies (22, 23) and found low correlation even when analyzing the effects of identical gRNAs targeted the same position. No correlation was observed between the gRNA effects in iPSC cells from Chen et al. (22) and those observed in the present study. However, we detected weak correlations between K562 cells from Chen et al. and cells used in the current study, with a slightly elevated correlation for the K562 and HAP1 pair, which may be attributed to their shared leukemic origin (Supplement Figures S5B–G). We observed weak correlations in 9 pairs produced by comparing three cell lines in the present study and cells from Schlesinger et al (25) (Supplement Figure S7). These findings reinforce the concept that uORF function is highly cell-dependent, as different screening approaches yield varying results, emphasizing the critical importance of the cellular context in assessing the uORF functionality.

Among a pool of significant hits in three cell lines, the majority were novel uORFs. However, we also identified several previously studied uORFs, including those in *ASDSD1* (63), *PPP1R15B* (94, 95), *TLNRD1* (96), *EIF2D* (97), *MIEF1* (65, 98). These uORFs can be classified into two major categories: microprotein-coding and regulatory.

The ASDURF microprotein (*ASDSD1* uORF) is linked to the PAQosome co-chaperone function (63) and essential in medulloblastoma (24). Our findings suggest that ASDURF may also be critically important in other cancers under specific conditions (Supplement Figure S9A). The MIURF microprotein (*MIEF1* uORF) has been implicated in mitochondrial translation, yet it was not previously recognized as essential. Our results indicate that its function is may be more crucial than was previously assumed. Other uORFs in *PPP1R15B*, *TLNRD1*, and *EIF2D* have been associated with translational control, suggesting their important regulatory role in cell proliferation depending on the cellular context.

We accessed features that distinguish essential uORFs from non-essential ones. Surprisingly, we did not find nucleotide conservation or protein coding potential to be defining features of essential uORFs (Supplement Figures S8O, S9E). This may be due to their translation rather than their products being under selection. It also possible that their products became evolutionary important relatively recently. Thus, we leveraged another approach for discovering microprotein-coding uORF. Comparison with DepMap gene essentiality data supports the idea that many uORFs affect cell proliferation independently of their CDS (Supplement Figures S2I, S3I, S4I), suggesting coding for essential microprotein.

To validate this hypothesis, we selected the *TRAM1* gene, which has a conserved, potentially coding uORF for further investigation (Supplement Figures S10A–B). Interestingly, while DepMap data indicated that *TRAM1* is non-essential in all tested cell lines, our testing in HAP1 cells showed significant gene essentiality when targeting the *TRAM1* CDS (Supplement Figures S10C–D), contradicting DepMap data. Of note, HAP1 cells have near zero effect for *TRAM1* according to DepMap (Supplement Figure S2I). Additionally, mutations in the first exon were even more critical, likely due to the complete disruption of functional protein production (Supplement Figures S10C). Thus, our results strongly suggest that screening validation using individual gRNAs is very important step in CRISPR-Cas9 screenings, as high throughput experiments are prone to artefacts and random outliers.

In light of the findings regarding *TRAM1*, we proposed that regulatory activity of uORFs is the primary driver of the uORF selection in the screening. Inspection of 5’ UTR sequences of selected genes suggested that in many cases, indel mutations lead to the uORF extension and overlap with the CDS. Such arrangement would likely result in the repression of the CDS translation due to the absence of translation reinitiation. If the encoded protein is important in a given cell-specific context, then it will inhibit cell proliferation. Frameshift mutations in uORFs lead to an overlap of uORFs with the CDS, substantially downregulate the CDS translation, and disrupts translational control of the *SLC25A39* and *YIF1A* genes (Figure 4A–B). Notably, our analysis revealed that essential uORFs tend to be shorter in length and are located within shorter 5′ UTRs (Supplement Figures S8F–G). This observation indicates that within short 5’ UTR, there are higher chances of an indel leading to the uORF overlap with the CDS and its translational repression.

We artificially introduced frameshifts in uORFs during the screening and in luciferase reporter experiments to disrupt the gene translation, however many known nucleotide polymorphisms that are present in 5’ UTRs. To explore the potential disease implications, we analyzed short nucleotide variants within uORFs in the ClinVar and dbSNP databases. Although ClinVar data on 5′ UTRs are limited, we identified numerous uORF-altering variants in dbSNP, including insertions, deletions, and single-nucleotide polymorphisms that may disrupt uORF regulatory functions (Figure 4D–E, Supplement Table S5). The indels lead to frameshifts in uORFs and can possibly repress the CDS translation and affect proliferation of specific human cells. Moreover, it was previously established that variants creating new upstream start codons are under strong negative selection (108), highlighting high relevance of uORF influence on gene expression. We anticipate that many more such variants will be discovered, as clinical studies traditionally prioritize the investigation of CDS regions. Our results also suggest that uORF-bearing genes with short 5’ UTR are more clinically relevant in this aspect, due to a higher likelihood that an indel will extend the uORF and repress the CDS translation.

In this study, we identified numerous novel uORFs that are essential for cell proliferation. Since only a small proportion of uORFs exhibit clear protein conservation, their essentiality is presumably driven by regulatory roles rather than peptide function. This raises an important question: if an uORF regulates an essential gene *in cis*, but does not encode a functional micropeptide, should it be considered as an important genetic element and annotated accordingly? Alternatively, should we continue to view the whole system as a single gene with complex translational regulation for the sake of simplicity? We propose that uORFs should be regarded as an essential genetic element if they have a significant impact on the CDS translation. We believe that Ribosome Decision Graphs can be applied for this to demonstrate the complex translation process in such cases (8).

The present findings provide a valuable set of uORF candidates for further functional analysis and highlight the need for further investigation into their mechanisms of action and translational control in specific cellular contexts. While we cannot rule out the possibility that other elements in the 5’ UTR of studied genes contribute to their expression and cell proliferation, such 5’ UTRs remain a subject of great interest for future studies. Unfortunately, some highly conserved uORFs could not be targeted due to PAM-motif constraints. Future research can leverage alternative CRISPR systems to edit these uORFs and assess their function in various genetic contexts.

In summary, we identified 155 uORFs that are essential for cell proliferation. The majority of these show strong cell type specificity and exert their function via *cis*-regulatory mechanisms that modulate translation of the downstream CDS rather than by encoding conserved micropeptides. Importantly, our analysis suggests that genes with short uORFs or located within short 5′ UTRs are particularly vulnerable to frameshift mutations that extend uORFs, leading to the CDS translational repression and reduced cell proliferation in case of essential genes. Numerous naturally occurring variants may disrupt the function of essential uORFs and contribute to human disease, underscoring the clinical relevance of these elements. Taken together, these findings expand our understanding of the regulatory landscape of the 5’ UTRs, thus highlighting uORFs as critical cell- and context-dependent modulators of gene expression. Our results represent a valuable resource for future studies that aim to dissect the post-transcriptional regulatory code and its implications in disease and therapeutic targeting.

## Data availability

The data underlying this article are available at NAR online.

## Supplementary data

Supplement Table S1 – uORF library

Supplement Table S2 – list of positive control genes Supplement Table S3 – gRNA library

Supplement Table S4 – screening results for HAP1, HEK293T, A549

Supplement Table S5 – nucleotide variants affecting uORFs from ClinVar and dbSNP

Supplement Table S6 – list of oligonucleotides used in the study

## Author contributions

Nikita M Shepelev: Conceptualization, Methodology, Investigation, Formal analysis, Validation, Visualization, Data curation, Writing – original draft. Elizaveta A Razumova: Formal analysis, Validation, Investigation. Alexandr I Lavrov: Formal analysis, Validation, Investigation. Stephen J Kiniry: Data curation, Methodology. Alexandr M Makaryuk: Investigation, Validation. Renata D Bibisheva: Investigation, Validation. Olga A Dontsova: Supervision, Writing – review & editing. Pavel V Baranov: Conceptualization, Resources, Writing – review & editing. Maria P Rubtsova: Conceptualization, Funding acquisition, Methodology, Supervision, Writing – review & editing.

## Acknowledgements

We are grateful to the Moscow State University Development Program NAUKA for providing access to the flow cytometers LongCyte (Challenbio) and SinoCyte (BioSino), the SeqStudio Genetic Analyzer (Applied Biosystems), and the Victor X5 plate reader (Perkin Elmer).

