## Supplementary figures and images for "Translation of numerous upstream open reading frames rather than their products is essential for proliferation of human cells of distinct origin"

### Supplemental Figure 1

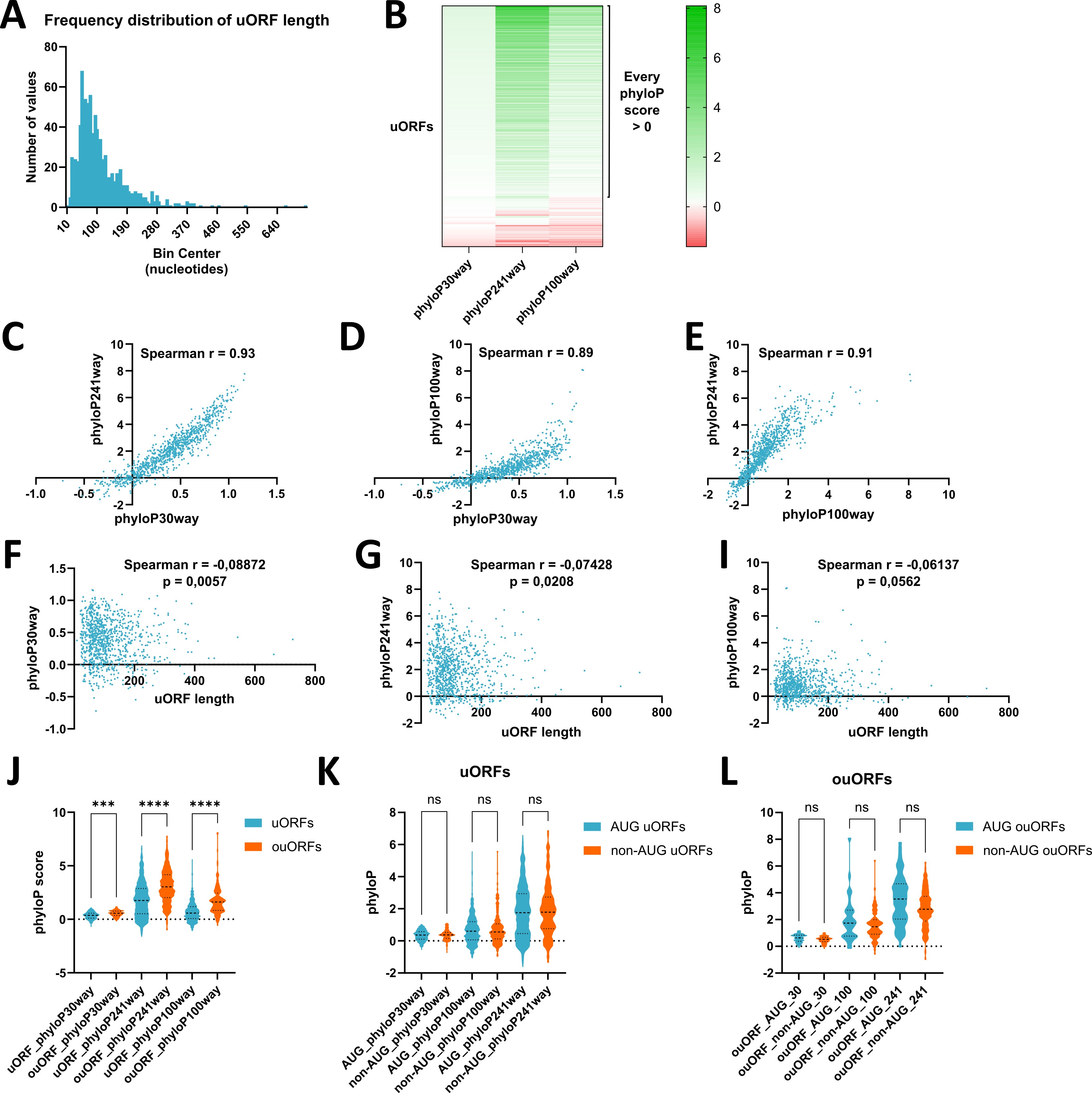

### Supplemental Figure 2

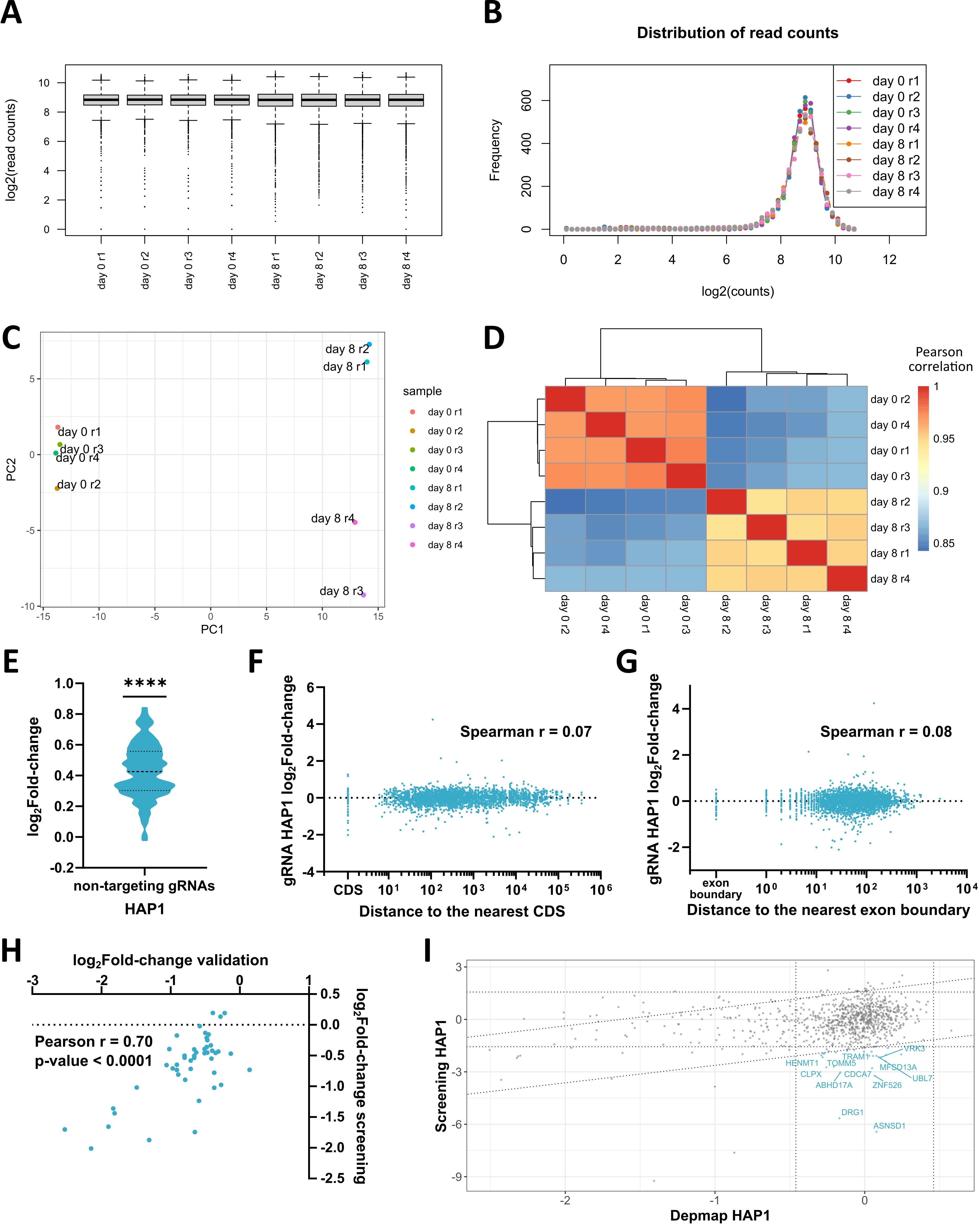

### Supplemental Figure 3

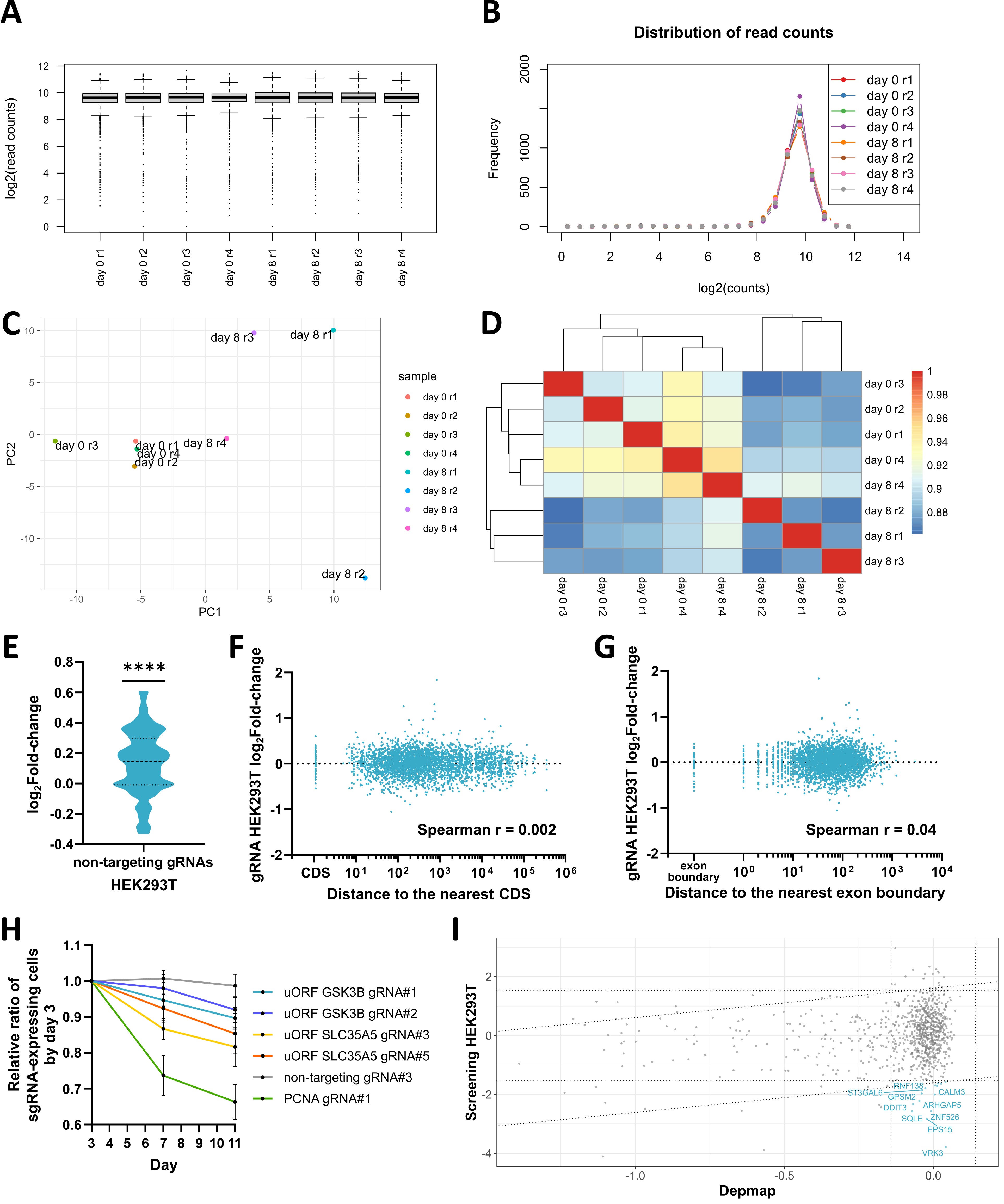

### Supplemental Figure 4

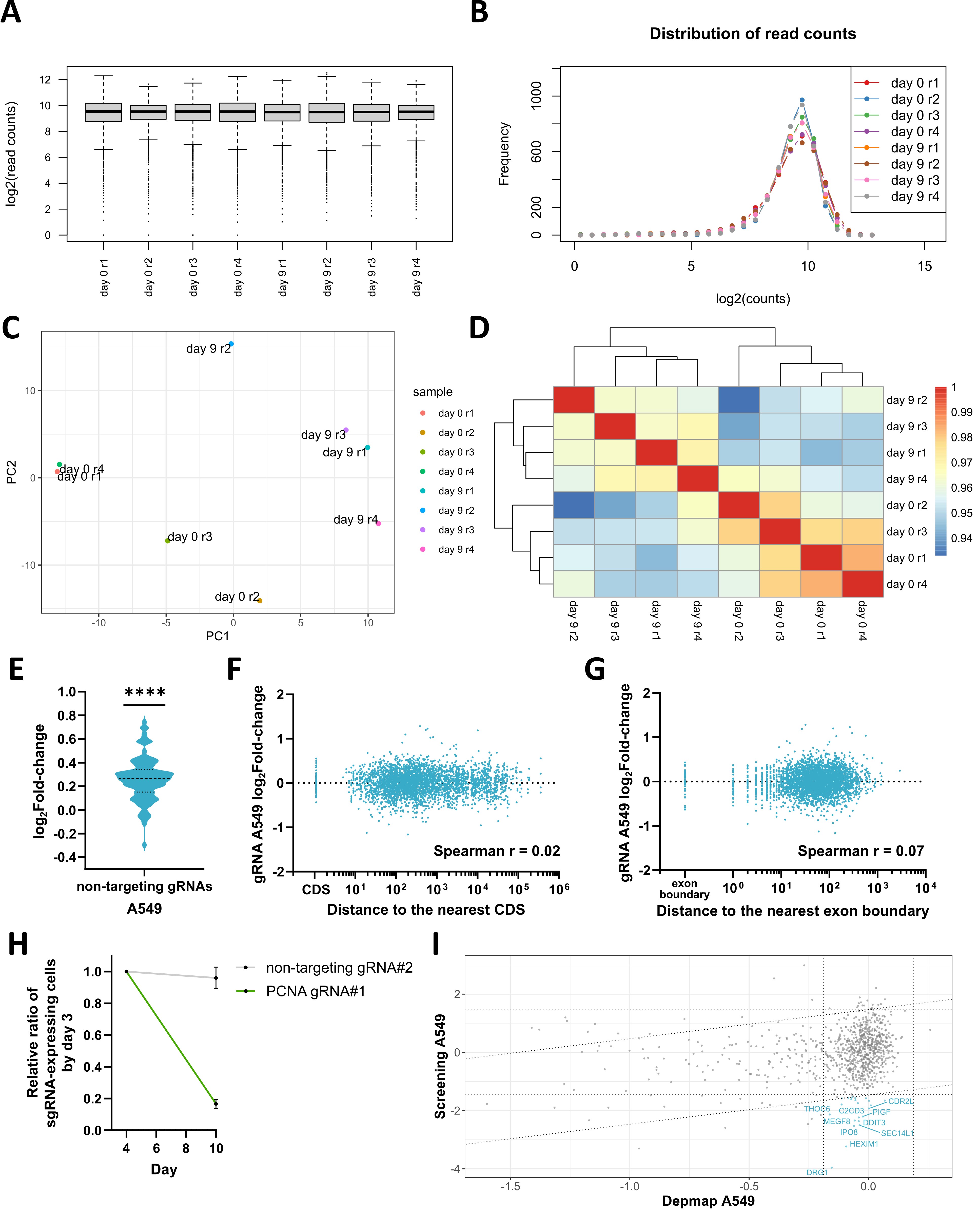

### Supplemental Figure 5

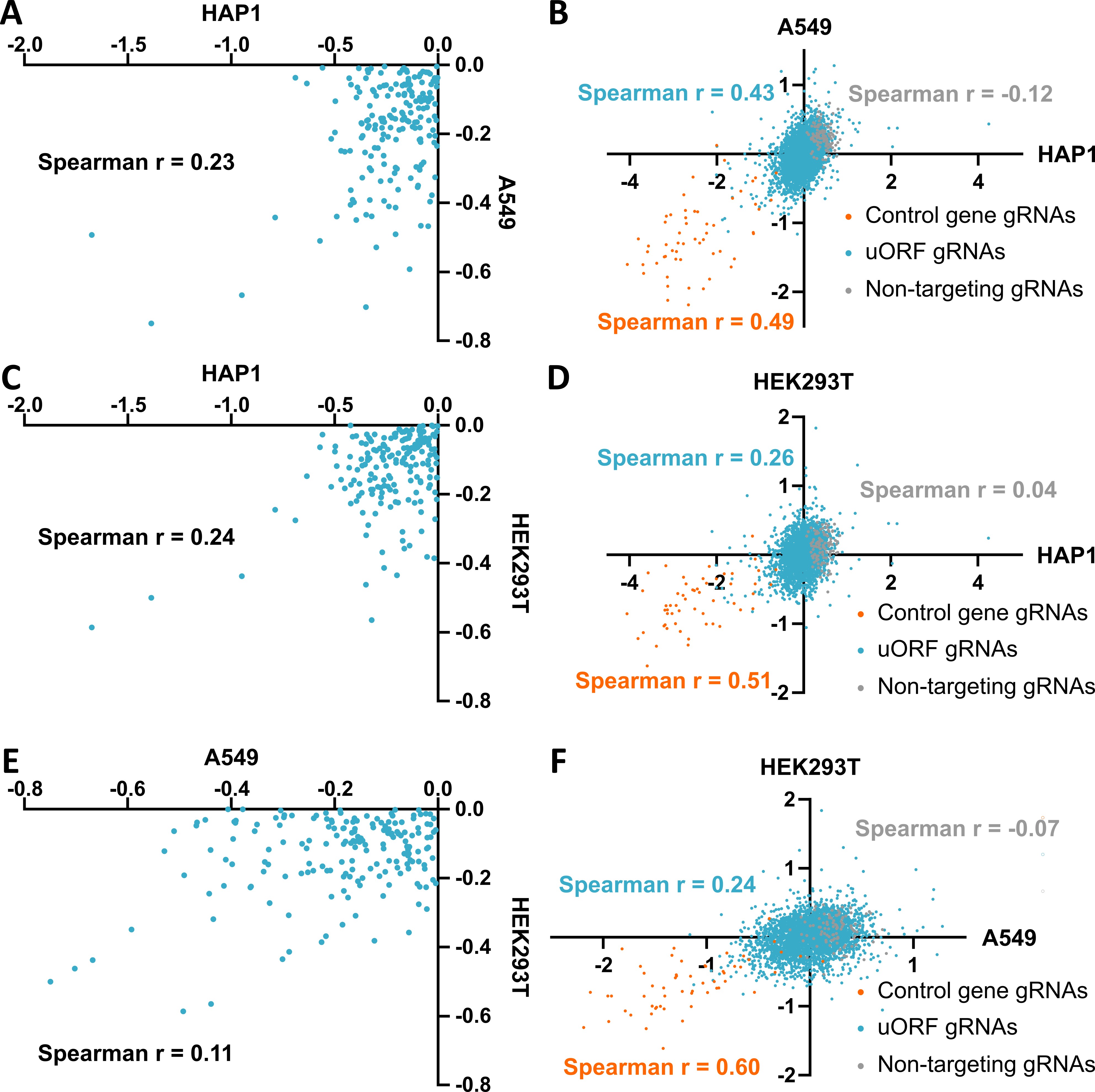

### Supplemental Figure 6

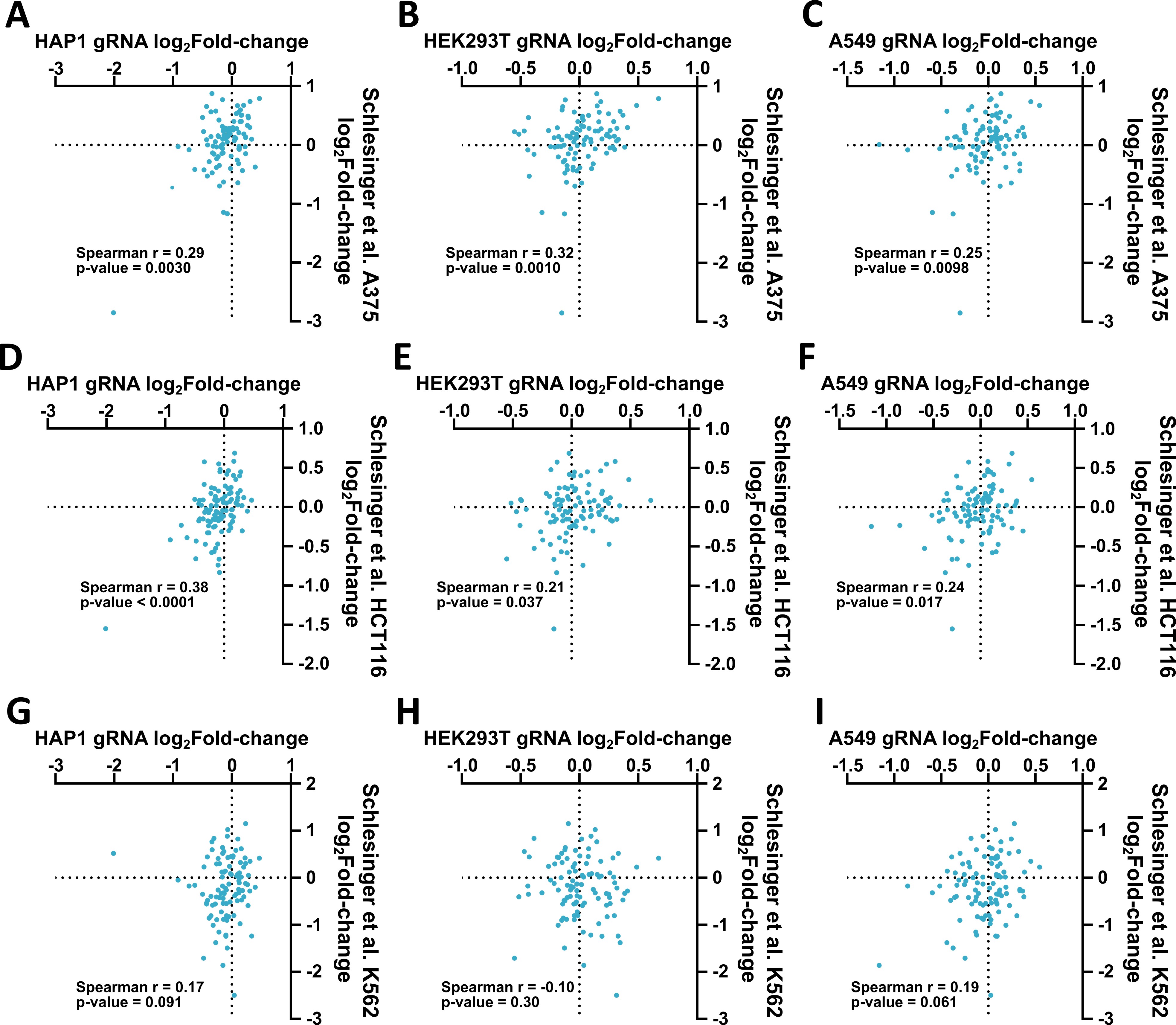

### Supplemental Figure 7

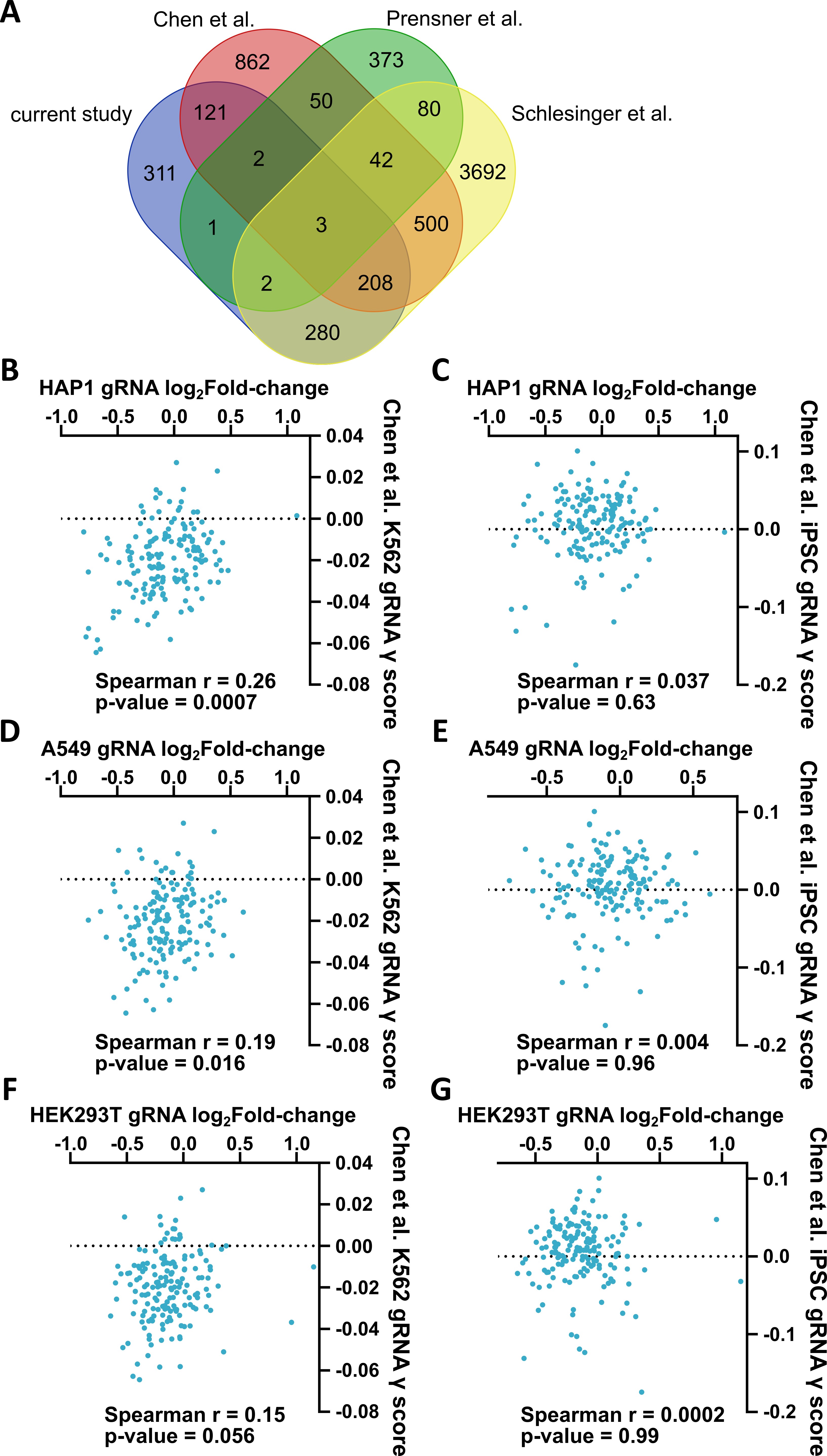

### Supplemental Figure 8

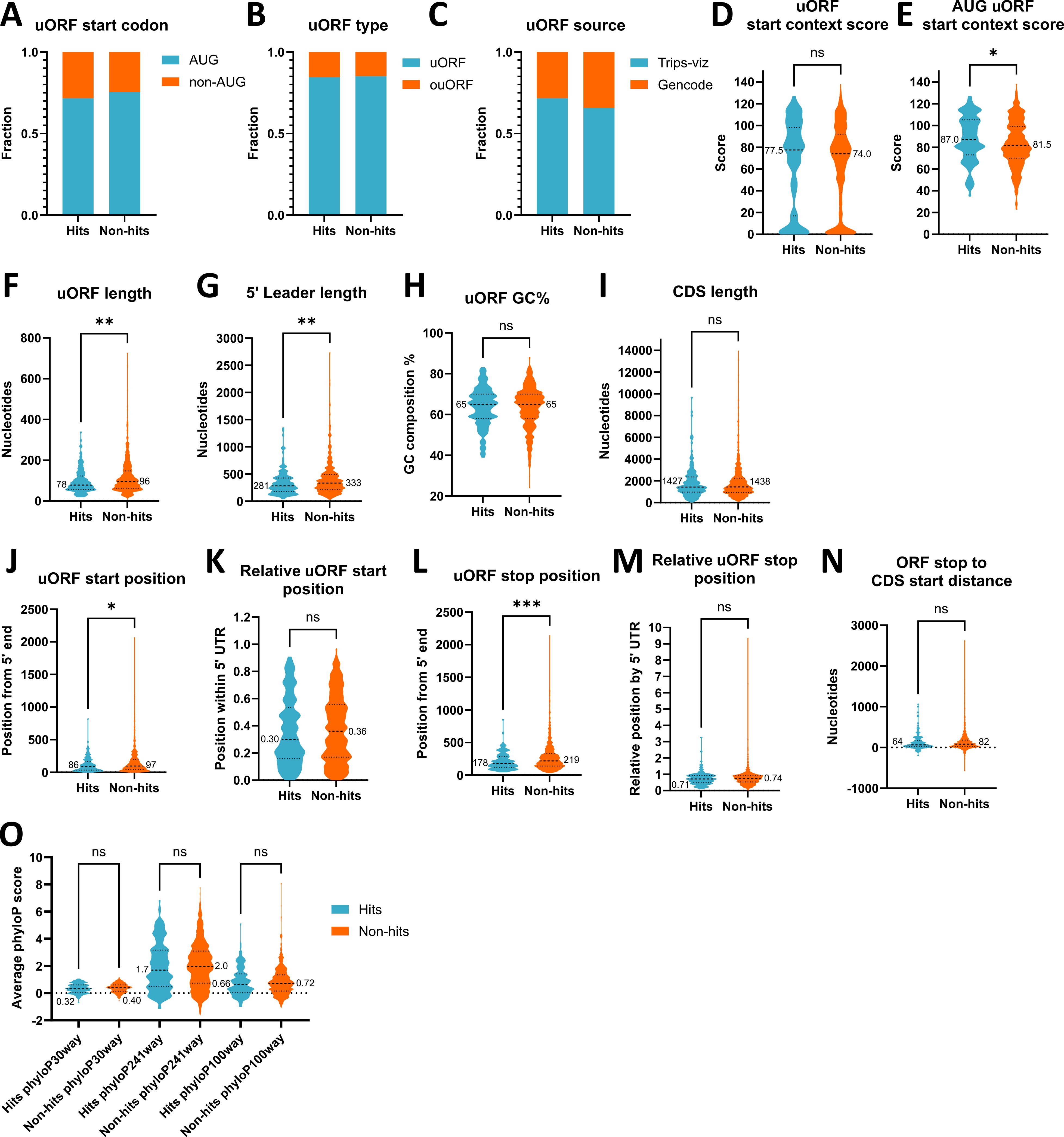

### Supplemental Figure 9

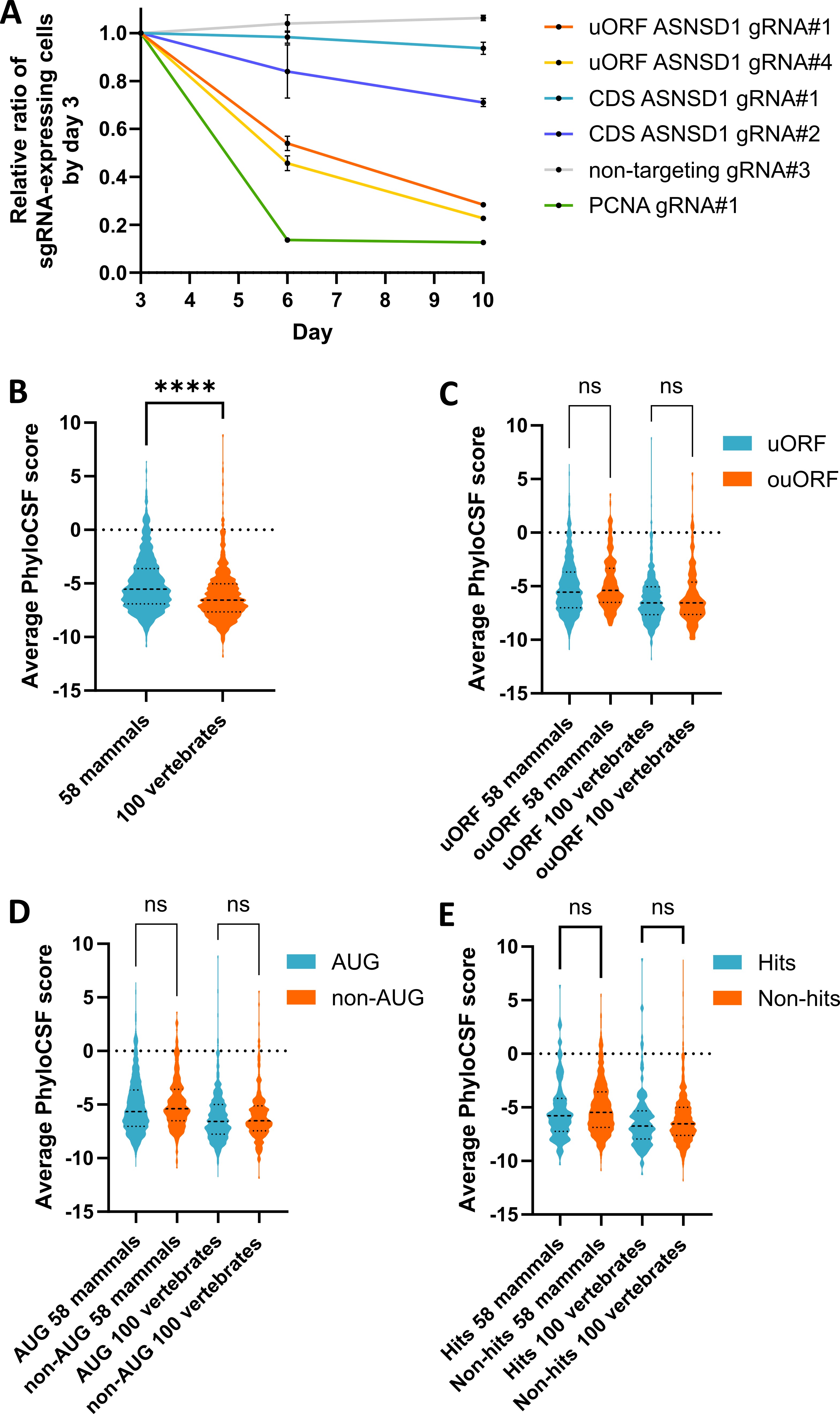

### Supplemental Figure 10

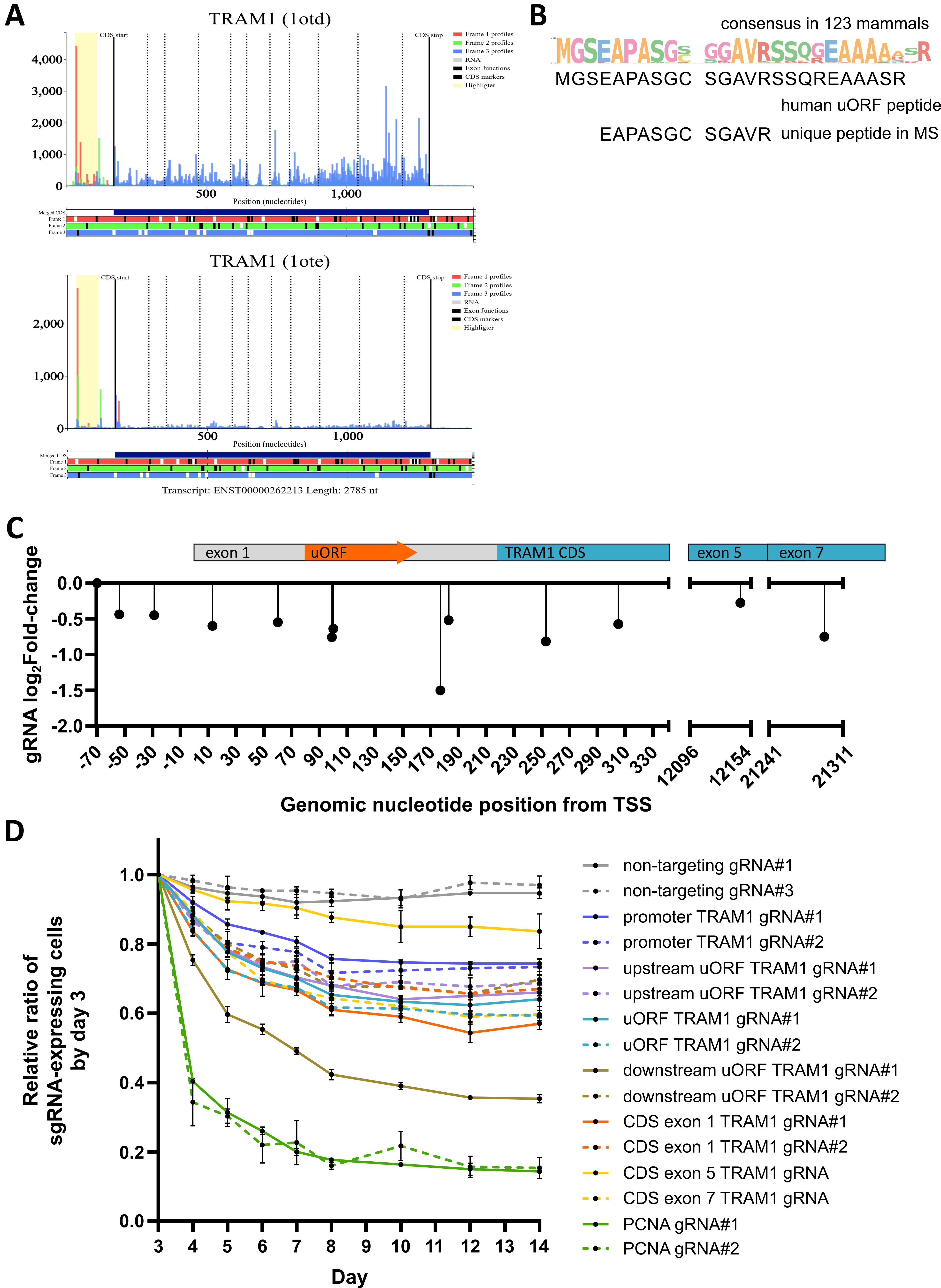
