## Supplemental Table 2 for "Translation of numerous upstream open reading frames rather than their products is essential for proliferation of human cells of distinct origin"

Supplement Table S2. The list of positive control genes.

| **Gene name** | **Full name** | **Cell function** |
| --- | --- | --- |
| CDC45 | Cell Division Cycle 45 | Initiation of DNA replication |
| EEF2 | Eukaryotic Translation Elongation Factor 2 | Translation elongation |
| EIF4A1 | Eukaryotic Translation Initiation Factor 4A1 | Translation initiation |
| EIF5 | Eukaryotic Translation Initiation Factor 5 | Translation initiation |
| HSPA9 | Heat Shock Protein Family A (Hsp70) Member 9 | Protein folding and maturation |
| MTOR | Mechanistic Target of Rapamycin Kinase | Regulation of cellular metabolism and growth |
| PCNA | Proliferating Cell Nuclear Antigen | DNA replication |
| POLR1B | RNA Polymerase I Subunit B | Transcription |
| PRPF19 | Pre-mRNA Processing Factor 19 | mRNA processing |
| PSMA6 | Proteasome 20S Subunit Alpha 6 | Protein degradation |
| SF3B3 | Splicing factor 3b subunit 3 | Splicing |
| TRMT112 | tRNA Methyltransferase Activator Subunit 11-2 | RNA and protein methylation |
| TUBB | Tubulin Beta Class I | Cytoskeleton |
| XPO1 | Exportin 1 | Transport |
